# A brain-to-text framework of decoding natural tonal sentences

**DOI:** 10.1101/2024.03.16.585337

**Authors:** Daohan Zhang, Zhenjie Wang, Youkun Qian, Zehao Zhao, Yan Liu, Xiaotao Hao, Wanxin Li, Shuo Lu, Honglin Zhu, Luyao Chen, Kunyu Xu, Yuanning Li, Junfeng Lu

## Abstract

Speech brain-computer interfaces (BCIs) directly translate brain activity into speech sound and text, yet decoding tonal languages like Mandarin Chinese poses a significant, unexplored challenge. Despite successful cases in non-tonal languages, the complexities of Mandarin, with its distinct syllabic structures and pivotal lexical information conveyed through tonal nuances, present challenges in BCI decoding. Here we designed a brain-to-text framework to decode Mandarin tonal sentences from invasive neural recordings. Our modular approach dissects speech onset, base syllables, and lexical tones, integrating them with contextual information through Bayesian likelihood and the Viterbi decoder. The results demonstrate accurate tone and syllable decoding under variances in continuous naturalistic speech production, surpassing previous intracranial Mandarin tonal syllable decoders in decoding accuracy. We also verified the robustness of our decoding framework and showed that the model hyperparameters can be generalized across participants of varied gender, age, education backgrounds, pronunciation behaviors, and coverage of electrodes. Our pilot study shed lights on the feasibility of more generalizable brain-to-text decoding of natural tonal sentences from patients with high heterogeneities.

## Introduction

Sentence is the basic language unit embodies our construal of representational meaning and interpersonal meaning, which constitutes the basis for daily communication^1^. Recent investigations have demonstrated the possibility of synthesis and decoding sentences in non-tonal languages ^2–11^ using intracranial neural recordings such as electrocorticography (ECoG) and Utah array. These studies have primarily relied on decoding the spatiotemporal neural patterns associated with articulatory movements—such as those of the lips, tongue, and larynx—in the ventral sensorimotor cortex during intended speech production. These advancements provided novel approaches for treating anarthria^12^ and enhanced the communication efficacy of speech brain-computer interfaces (BCIs)^13^.

However, decoding tonal sentences is still a largely-unexplored work. More than 60% of the languages in the world are tonal^14^, with approximately 2 billion people speaking tonal languages, including most Sino-Tibetan languages and the entire Tai-Kadai family^15^. Pitch in these languages is used to distinguish lexical and grammatical meaning^13^. While prior research has investigated decoding stereotypical instances of lexical tones from neural activity for monosyllabic speech^13^, decoding continuous tonal sentences is still a challenging issue. Unlike the relatively stable acoustic cues in canonical forms, natural speech introduces substantial variability in tone components. According to Fujisaki model, these components, typically represented by base frequency (F0) contours, encompass base frequency, phrase variations, and accent components^16,17^. Furthermore, the influence of tone sandhi—alterations in a morpheme’s tone due to syntactic context—adds an additional layer of complexity^18^. Altogether, these variances make decoding tonal sentences more complicated than both non-tonal sentences and isolated tonal syllables.

Besides, existing invasive language BCI usually reported success in individual cases, usually with intense hyperparameter optimizing^5,9,10^. Few studies have tested the replication or generalization of the decoding framework. It remains uncertain whether the same set of model design hyperparameters—such as the number of convolution and recurrent blocks, hidden variables per layer, and onset detection thresholds—will generalize across different subjects. Consequently, verifying the generalizability of these published frameworks, especially when applied to patients exhibiting high heterogeneity, remains elusive. In clinical settings, different patients need to restore speech function via speech BCI have varied pronunciation behaviors, which is especially obvious among tonal languages such as Mandarin Chinese. Mandarin speakers, influenced by regional dialects such as Northern, Jianghuai, and subgroups like Wu-Tai, Shanghainese-influenced, Northeastern Mandarin, demonstrate distinct tonal and syllabic variations^19^. Though sharing the same written form of Chinese word, speakers of these mandarin branches have distinct pronunciation behaviors on both tones and syllables. While these distinct pronunciation behaviors or preferences do not affect speakers’ daily communication, they add significant challenges to the decoding task. In addition to the variance in pronunciation behaviors, the inherent variations in language-functional cortical areas also contribute to the inter-subject variances. It is impossible for invasive devices to collect identical neural signals from different patients even if the placement of invasive electrodes is stringently controlled. Whether and to what extent these bias from both language behaviors and varied neural signals will affect the robustness of language decoding framework remain unknown. This uncertainty challenges the eventual generalization of speech BCI across patients with high heterogeneity.

In this study, we aim to decode Mandarin tonal sentences from invasive neural recordings using high-density ECoG. Targeting functionally separate neural populations within speech-related brain regions, we tailored distinct neural network modules which detected the onsets of the utterance of each individual Chinese character and then decode tone labels and syllable labels in parallel, reflecting the inherent parallel coding previously observed in tonal language articulation^13,20–22^. Subsequently, a language model was used to calculate the Bayesian likelihood of the entire sentence from the probability distribution of tonal syllable sequences, integrating the contextual and prior information (**Fig.1**). Moreover, we established a versatile framework using a set of standardized hyperparameters, eliminating additional needs for hyperparameter optimization, and assessed its potential for generalization by testing across multiple patients. This approach decoded Mandarin tonal sentences across diverse patient profiles without specific hyperparameter adjustments.

**Fig 1.**
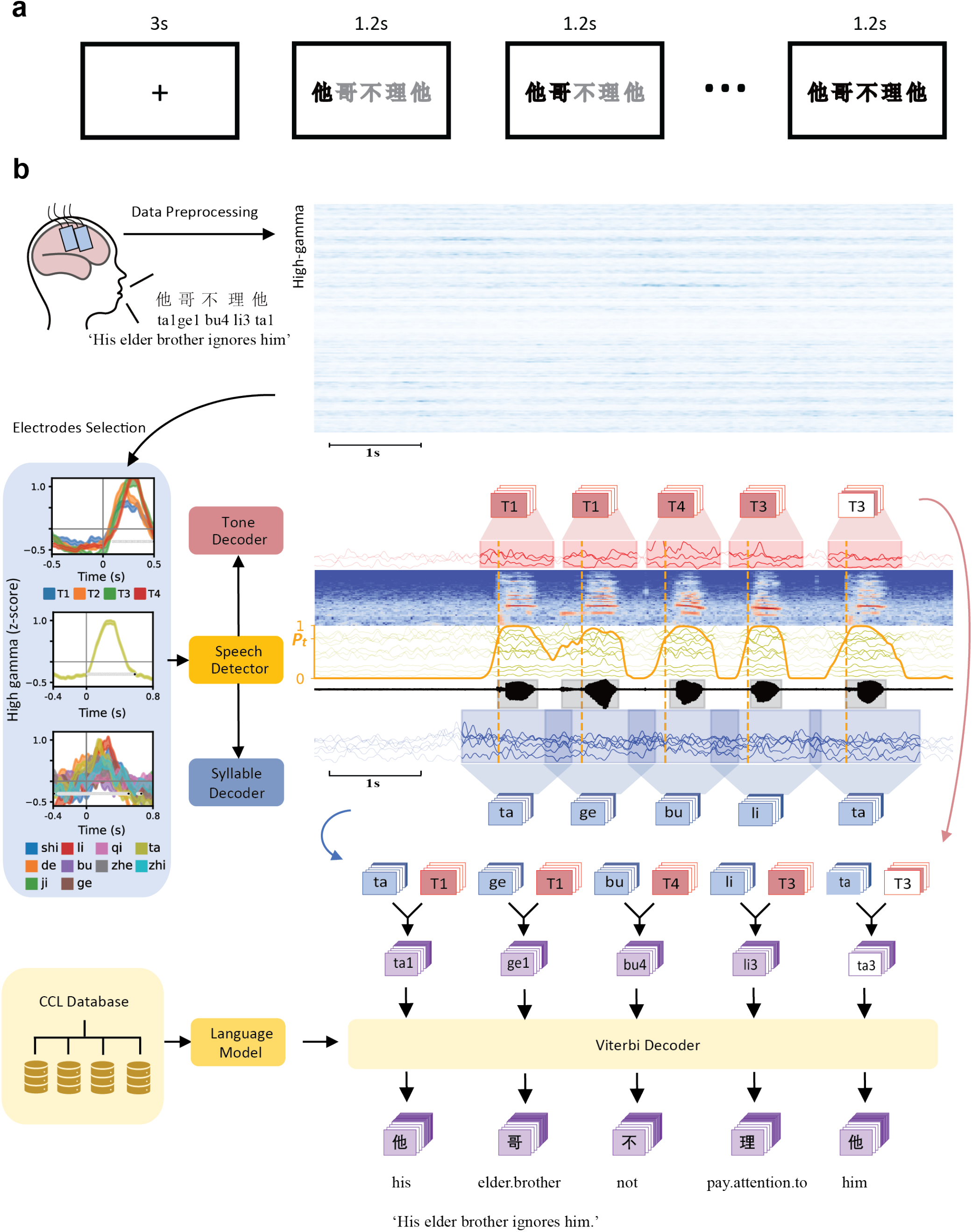
Schematic overview of the brain-to-text decoder for natural speech of a tonal language. a. Schematic overview of the tonal speech production task. Each participant was guided by a visual cue to produce one of the ten sentences. Each trial began with a fixation cross at the center of the screen for 3 seconds, and the sentence was shown in the middle of the screen in grey text. With each word turning black for 1.2 seconds consecutively from the beginning to the end of the sentence, the participant was instructed to pronounce the sentence following these go cues, the pace of the speech was not strictly aligned with the visual cues. b. During the experiment, neural activity from speech-related cortex was recorded using implanted high-density electrocorticography array when the participant was instructed to read sentences consisted of words from a predefined vocabulary set of 40 words. The preprocessed neural signals from responsive electrodes were sampled by sliding window and was sent to the speech detection module (speech detector) to detect the onsets of words. The peri-onset neural activity within a fixed time window was used to compute the base syllable probability (across 10 possible syllables) and lexical tone probability (across 4 tones) of the word via a tone decoder and a syllable decoder respectively. A Viterbi decoding algorithm used these probabilities in conjunction with word-sequence probabilities from a pre-trained natural language model to decode the most likely sentence given the current neural activity.

## Results

In this study, we recorded the neural activity of five native Mandarin-speaking participants who underwent awake surgery to treat brain tumors. Each participant was guided by sequential visual cues to produce 10 sentences consisted of 5-8 Chinese characters from a corpus of 40 Chinese characters with varied tones **(Fig. 1a)**. Among 5 participants, PA1-4 completed the speech production task using normal articulating, while PA5 completed the speech production task using whispering. Participants’ brain activity was recorded by temporally placed high-density ECoG grids. Subsequently, we assessed the efficacy of our proposed brain-to-text speech decoder across these 5 participants.

### Decoder overview

Our brain-to-text decoder comprises interconnected modules: a speech detector, tone decoder, syllable decoder, and a language model, functioning in a sequential stream. To decode Mandarin tonal sentences from high-density ECoG neural recordings, our brain-to-text decoder starts with a speech detector, which takes in sequence of neural activity recordings of pre-selected speech-responsive electrodes and predicts whether each timepoint belongs to speech production or resting-state. The output of the speech detector was used to identify speech production epochs, where each epoch corresponding to the time interval of one Chinese character. Based on the decoded speech production epochs, our tone and syllable decoders then decode tone label and syllable label from neural signals of pre-selected tone and syllable discriminative electrodes during each epoch.

For PA1 to PA4, we identified 108,144,199, and129 speech responsive electrodes, 30,28,68, and 75 syllable discriminative electrodes and 10,12,21, and 48 tone discriminative electrodes, respectively. For PA5, we found 118 speech responsive electrodes, 47 syllable discriminative electrodes and 14 tone discriminative electrodes. The coverage and overlapping relationship of these electrodes were shown in **Fig. 2a-e**.

**Fig 2.**
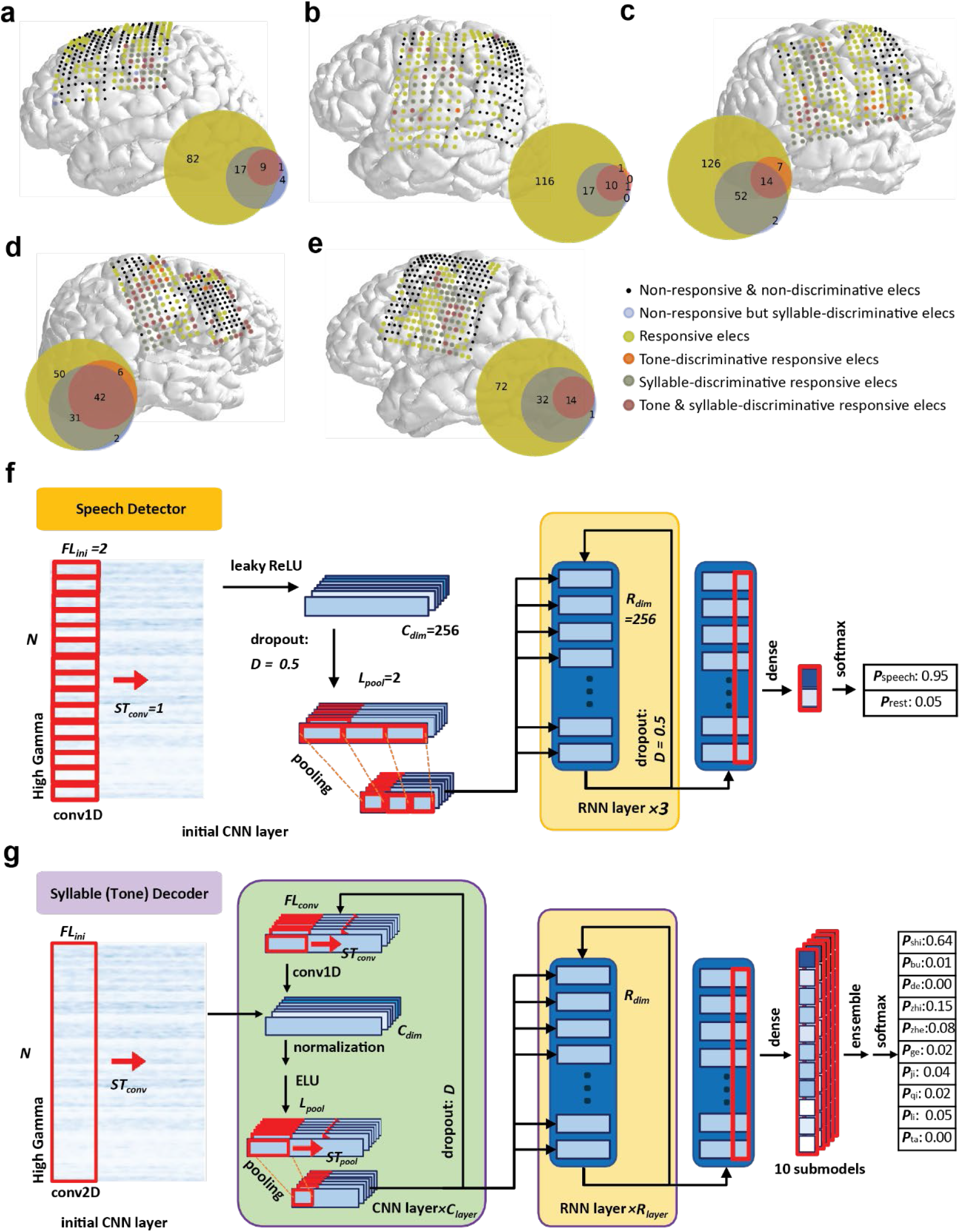
Electrode coverage, category and decoding model schematics. a-e. Anatomical reconstructions of PA1-PA5 (from a to e). The locations of the ECoG electrodes were plotted with colored discs. The colors indicated the electrode categories. Yellow: responsive electrodes; red: tone-discriminative electrodes; blue: syllable-discriminative electrodes. Electrodes with combined feature were plotted with mixed colors, nonresponsive electrodes were plotted as small black dot. Venn diagrams showed the number of electrodes in each category for each participant. f. Speech detection model schematic. Predefined hyperparameters in ANN and their values were shown in italic. g. Syllable classification model schematic. CNN unit shown in green frame while RNN unit shown in yellow. All of the undefined hyperparameters were shown in Italic, which can be roughly divided into time-dimension-related group and size-related group. Former group includes initial convolution filter length (***FL_ini_***), stride of convolution (***ST_conv_***), filter length of following convolutional blocks (***FL_conv_***), max-pooling kernel length (***L_pool_***) and max-pooling stride (***ST_pool_***). Later one including number of sequential convolutional blocks (***C_layer_***), number of layers of RNN (***R_layer_***), number of filters in each convolutional process (***C_dim_***), number of dimensions in each RNN process (***R_dim_***), and dropout value (***D***)) The tone classification model shared the same architecture with syllable classification model but different hyperparameters.

For onset predicting, the neural activity across all speech-responsive electrodes was processed time point by time point by an artificial neural network (ANN) containing sequentially arranged Convolutional Neural Network (CNN) structure, a stack of Gated Recurrent Unit (GRU) layers and a single dense (fully connected) layer, capturing both forward and backward temporal dependencies in neural signals, which was designated for inference on dynamic temporal processes. The dense layer projected the latent dimensions of the last GRU layer into probability space for two event classes: speech and rest. **(Fig. 2f)** For simplify, all the hyperparameters in this step was fixed. After that, the curve of the predicted probabilities along the dimension of time was smoothed and binarized according to undefined smoothing window (***S***) and probability threshold (***P_t_***), the onset of each utterance was then predicted if a lasting silence states (***T_off_***) before the onset and a lasting speech state (***T_on_***) after it, while slightly errors labels within an undefined error permissive rate (***EPR***) was also allowed **(Fig. 3b)**. These five undefined hyperparameters need further tuning through grid search for each participant.

**Fig 3.**
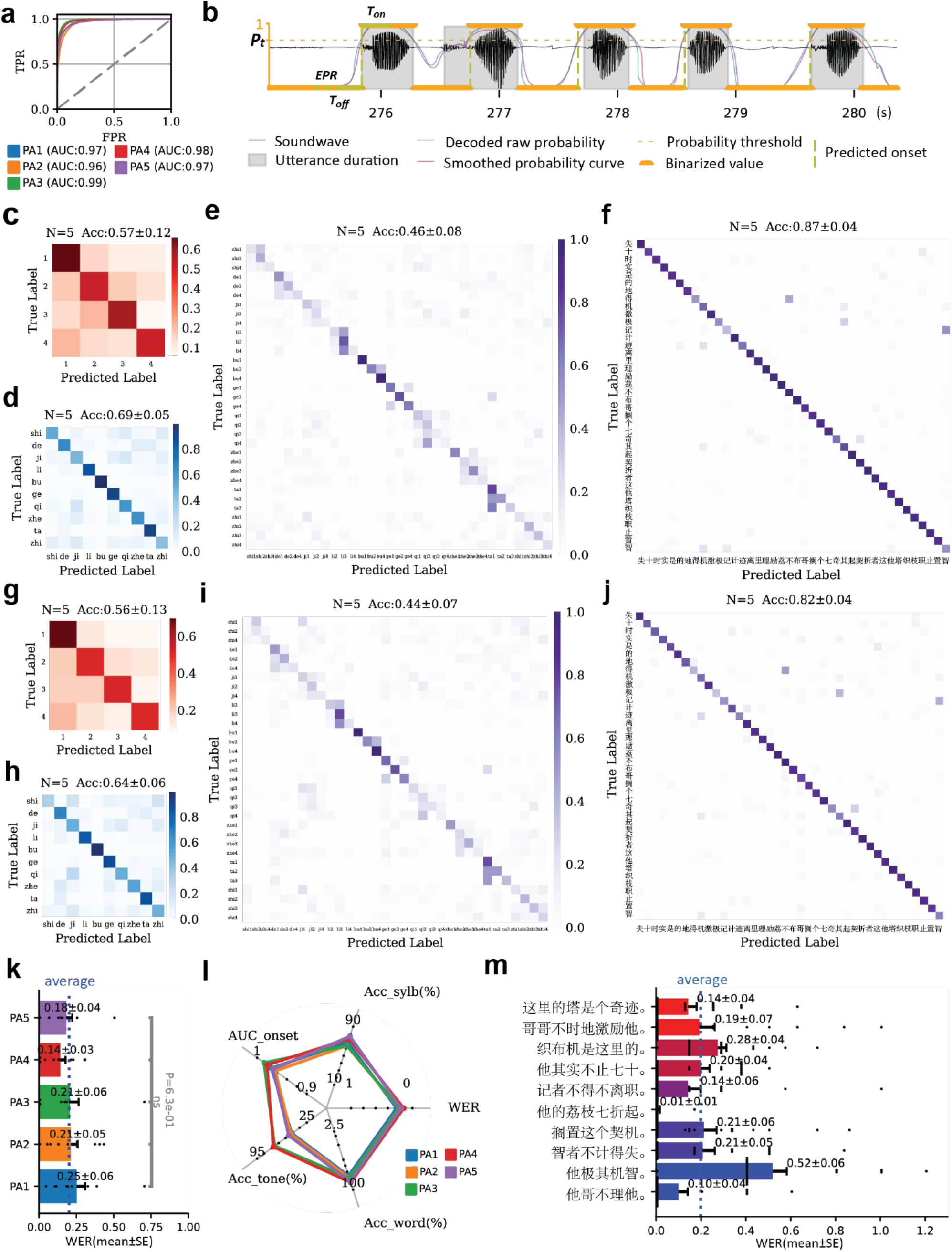
Evaluation of the overall neural-to-text decoding performance of the decoder. a. Receiver Operating Characteristic (ROC) curves and corresponding area under curve (AUC) of speech detectors in each participant. b. One example trial, in which the participant produced the sentence “他哥不理他” (His elder brother ignores him). The raw speech sound waveform was plotted in black. The time course of the predicted speech probability was plotted blue, while speech probability after smoothing and binarized was plotted red and orange. Finally, the detected speech event onsets from the neural decoder was plotted as yellow dotted vertical lines. All of the undefined hyperparameters were shown in Italic. c-f. Confusion matrices of c) the tone labels, d) the syllable labels, e) the tonal syllables, and f) the words (with language model), evaluated on the test set, using manually aligned actual speech onsets. g-j. Confusion matrices of g) the tone labels, h) the syllable labels, i) the tonal syllables, and j) the words (with language model), evaluated on the test set, using decoded speech onsets from the onset decoder. k. The averaged word error rate (WER) of decoded sentences in 5 participants, mean [±sem], the vertical dotted line indicated the overall averaged performance across all 5 participants. l. Performances of the speech detector, the tone decoder, the syllable decoder and the overall performances of each participant (in the same color keys as panel **k**), shown by AUC of speech-silent classifier (AUC_onset), tone accuracy (Acc_tone), syllable accuracy (Acc_sylb), word accuracy (Acc_word) and WER respectively. m. The averaged WER of each individual sentence across all 5 participants, mean [±SE], dotted line indicated the average performance across all sentences.

As for tone and syllable decoder, a 1.2-second time window for syllable and 0.8-second time window for tone of high gamma activity was processed by an ensemble of 5 pairs (10 total) ensemble ANN models. ANNs in tone and syllable decoder share the same architecture but different in their own hyperparameters. Within each ANN, the high gamma activity was processed by an initial convolution with initial filter length (***FL_ini_***), stride of convolution (***ST_conv_***). This initial layer was followed by undefined number (***C_layer_***) of CNN unit^2^. Each CNN unit constituted of a temporal convolution with undefined kernel length (***FL_conv_***), aforementioned stride length and dimension (***C_dim_***), a batch normalization, ELU activating function^23^, a dropout layer with dropout rate (***D***) and maxpooling layer with max-pooling kernel length (***L_pool_***) and max-pooling stride (***ST_pool_***). After that, data was processed by undefined number (***R_layer_***) of stacked bidirectional gated recurrent unit (GRU) layers with undefined number of dimension (***R_dim_***) ^24^. A dense layer projected the final GRU layer into probability of syllable and tone of each of the words from the 10-syllable set or 4-tone classes. Finally, we averaged these probability distributions from ensembled ANN models to get the predicted syllable and tone probabilities. **(Fig. 2g)** It is worth noting that all the aforementioned undefined hyperparameters required further tuning for each participant.

Subsequently, a language model integrates these decoded tone and syllable labels, along with prior information of their transitional probabilities, to compute the Bayesian likelihood of entire word sequences.

### Independent performance of speech detector, tone decoder and syllable decoder

First, we evaluated the performance of each individual decoder module. Since these decoder modules work in sequential order, the performance of the tone and syllable decoders would rely on the output of the speech detector. To evaluate the independent performance of these decoder modules, we first calculate the decoding accuracies of syllable, tone, tonal syllable, and Chinese characters on manually aligned speech onsets rather than onsets predicted by the speech detector.

Each of the participants completed 158 to 160 sentences in the speech production task. Using nested cross-validation, we trained the brain-to-text decoder for each participant and evaluated the decoding performance. Among five participants, our speech detector reached an area under curve (AUC) of 0.96 to 0.99 **(Fig. 3a)**. The tone decoder reached an accuracy of 57%±12% (mean [± SD] classification accuracy across the 4 target tones in 5 participants, chance 25%) on manually aligned neural signals **(Fig. 3c)**. Syllable decoder reached an accuracy of 69%±5% (mean [±SD] classification accuracy across the 10 target syllables in 5 participants, chance 10%) on manually aligned neural signals **(Fig. 3d)**. When multiply raw predicted probability of tone and syllable, we acquired raw decoding accuracy of all the predicted tonal-syllables, which reaches 46%±8% (mean [±SD] classification accuracy across the 40 tonal syllables on 5 participants, chance 3.0%) at the level of tonal syllable **(Fig. 3e)**. After applying language model to calculate corresponding Chinese words, the accuracy at the level of Chinese words reaches 87% ± 4% (mean [±SD] classification accuracy across the 40 Chinese words on 5 participants, chance 2.5%) **(Fig. 3f)**.

### Overall decoding performance of tonal-sentences

After evaluated the independent performance, we then tested the performance of all the modules when they were interconnected and worked in a stream. Aligned with speech onsets detected by speech detector, tone decoder achieved an accuracy of 56%±13% (mean [±SD], **Fig. 3g)**, while syllable decoder achieved an accuracy of 64%±6% (mean [±SD], **Fig. 3h)**. Accuracy at the level of raw tonal syllable reached 44%±7% (mean [±SD], **Fig. 3i)**, while the final accuracy at the level of Chinese words reaches 82%±4% (mean [±SD], **Fig. 3j)**. These results were consistent with the independent module performance, when actual speech onsets were manually aligned. We also calculated the word error rate (WER) of each decoded sentence. Decoding accuracy was consistent across participants (one way-ANOVA, *F(4, 45)* = 0.65, *p*=0.63), as the overall WER was 25%± 6%, 21% ± 5%, 21% ± 6%, 14% ± 3%, 18% ± 4% (mean [±SE]) from PA1 to PA5. For each individual sentence, WER ranged from 1% to 52%. 47.74% of all the sentences (95/199) were decoded correctly (WER=0) (**Fig. 3k**). There was no significant difference (Pearson’s correlation, r = −0.03, *P* = 0.61) between the decoding accuracy and the complexity (number of words or phrases) of the sentence (**Fig. 3m**), suggesting that our proposed method worked for both short and long sentences.

### Robustness of the speech decoder under tonal variance in natural speech

In natural speech, the actual pitch trajectories of lexical tones often deviate from their canonical forms, due to accent, emotions, and other contextual effect such as coarticulation and tone sandhi^25^. For example, all the patient articulated the tone of “不” in “不计得失” with a pitch trajectory more similar to tone 2 rather than tone 4 in single syllable form or in other phrases such as “不得不”, due to the rule of tone sandhi (Fig 4a). Even the same tonal syllable with different phonological context would result in different pitch trajectories (Fig 4b). This suggests that the underlying neural code commanding tone articulation may also encounter great variance. Therefore, the decoding algorithm should not only consider the stereotypical canonical monosyllable instances of lexical tones, but also able to account for such significant variance during natural speech and robustly decode lexical tones regardless of the variance.

**Fig 4.**
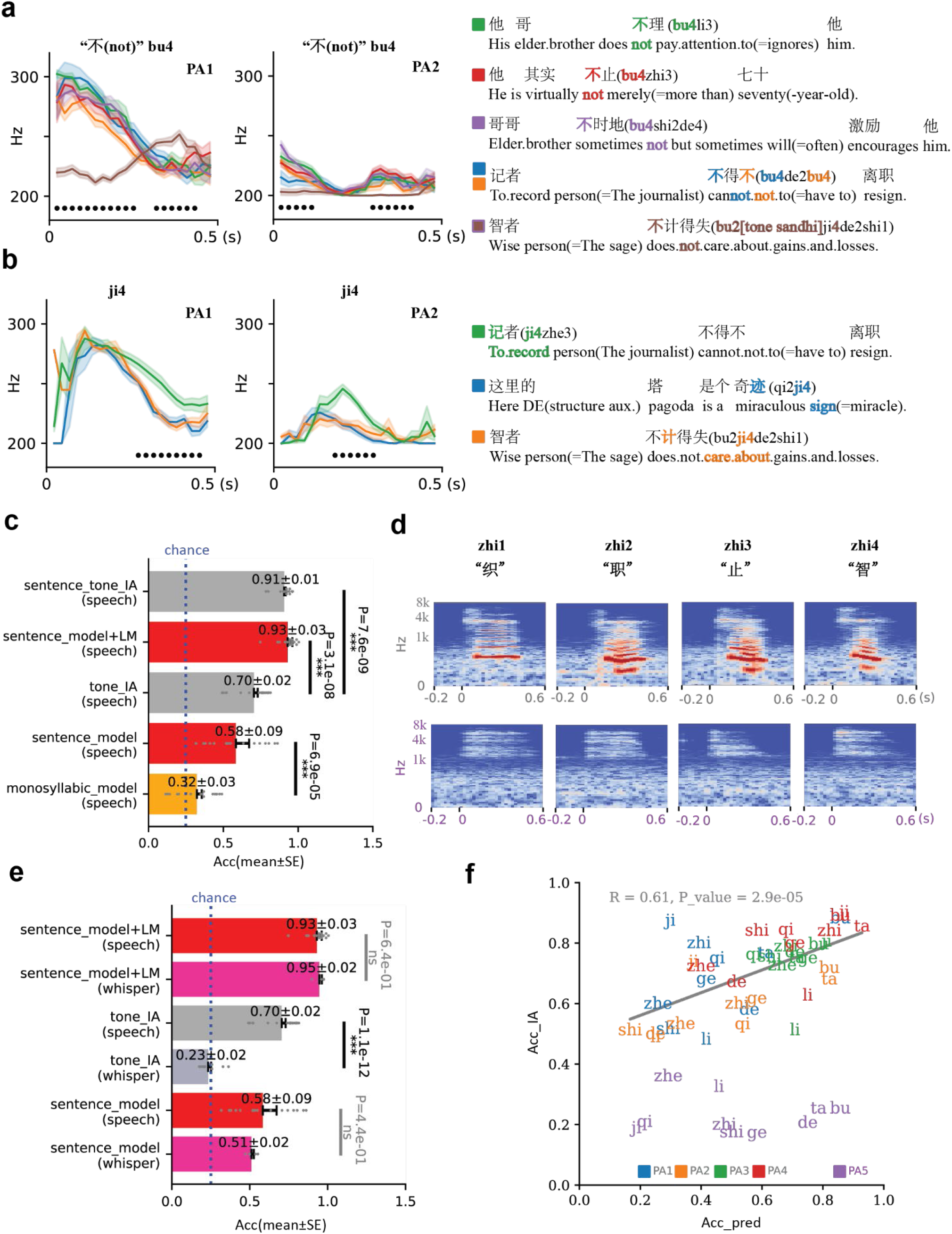
Evaluation of tonal decoding under natural contextual variances and whispering. **a-b)** Averaged pitch contours (mean [±SE]) of different Chinese characters of the same tonal syllables (a: bu4, also the same Chinese character “不” which represents negative meanings such as “not”, b: ji4), black dots indicated time points with significant mean difference (one-way ANOVA, *p* < 0.05). The text transcriptions were shown in the bottom with the specific words highlighted in the corresponding colors. Corresponding Chinese characters, *Pinyin*, and English translation of the choosen syllables were shown in the same color as the pitch contour at the right part of the figures. In the last sentence of the subplot **a**, the tone 4 of “ji” (also shown in color) leads to the change in the tone of previous “bu”, due to the rule of tone sandhi. **c)** Tone decoding performances (mean [±SE]) of previous monosyllabic neural decoding model (monosyllabic_model), our sentence-based neural decoding model without language model (sentence_model), our sentence-based neural decoding model with language model (speech_sentence+LM), the accuracy of tone intelligible analysis (tone IA score) by 20 volunteers using the corresponding speech audio syllable clips (tone_IA), and speech audio of each full sentences (speech_tone_IA). Blue dotted line indicated chance level (25%); *** *p* < 0.001, t-test, two-sided. **d)** The mel-spectrograms of participant (PA1) spoke aloud four tones of “zhi” (upper row), compared to the mel-spectrograms of participant (PA5) whispering four tones of “zhi” (bottom). **e)** Tone decoding performances using our proposed sentence decoder without language model (sentence_model) and with language model (sentence_model + LM), and the tone IA scores (mean [±SE]) evaluated on the speech and whisper participants. **f)** Scatterplot showing the correlation between of tone decoding accuracy (Acc_pred) and tone IA score (Acc_IA) of articulating participants (Pearson’s correlation, *r* = 0.61, *p* = 2.9×10–5). Whispering data shown in purple, which was not included in the linear regression due to randomly distributed tone IA scores.

First, we quantify the variance caused by tonal sentence context and tone sandhi behaviorally. We performed tone intelligibility assessment (IA) test through which native mandarin speaking participants listening to audio of each tonal Chinese word clipped from the natural tonal sentences and judging the tone. We find the accuracy of the tone IA test only reached 70%±2% (mean [±SE]), significantly lower (t-test, *t(38)* = 7.38, *p* = 7.6e-9) than behavioral performance under full natural context (tone IA accuracy 91%±1%, mean [±SE], Fig 4c). Therefore, using contextual information is important for listeners to overcome tonal variances in natural speech production.

Similarly, given only monosyllabic information, the tone decoder would perform suboptimal. To show this, we adopted a baseline monosyllable decoder model previously used in Liu et al^13^. The monosyllable decoder only took in the neural activity aligned to the current syllable utterance and did not consider contextual syllables. In our test dataset, the monosyllable baseline decoder model achieved an averaged tone decoding accuracy of 35%±3% (mean [±SE], Fig. 4**c**).

Finally, we tested the performance our proposed sentence decoder. We compared our tone-decoder (without the language model) with our previous published tone decoder designed for monosyllable^13^. Our framework (decoding accuracy 58%±9%, mean [±SE]) outperformed (t-test, *t(30)* = −4.62, *p* = 6.9×10-5) previous monosyllable tone-decoder (decoding accuracy 32%±3%, mean [±SE]) on tonal sentence decoding task, which suggests our framework is more robust on decoding tone components from natural language settings with multiple and unbalanced syllables. Furthermore, after introducing language model to our decoding framework, a tone decoding accuracy of 93%±3% (mean [±SE]) was achieved, which significantly exceed (t-test, *t(34)* = 7.12, *P* = 3.1×10-8, **Fig. 4c**) the accuracy of monosyllable tone IA results (70%±2%, mean [±SE]), and approximate the actual behavioral performance of native speakers under natural context (91%± 1%,). Hence, applying language model will largely eliminate the tone variance in natural language by introducing contextual relationships, which is a promising solution of decoding tone in natural tonal sentences accurately.

### Speech decoding during non-intelligible tonal speech production

To further evaluate the robustness of our proposed decoder under different speech production scenario, we tested if the decoder could decode from patient who did not produce intelligible speech. In particular, PA5 completed the speech production task using whispering and the produced tonal speech was largely non-intelligible. We first plot the Mel-spectrograms of Chinese words clipped from whispering natural tonal sentences, finding no obvious base frequency (F0) and consonant peak (**Fig. 4b**). Furthermore, the tone IA score of PA5 was 23%±2% (mean [±SE]), which was significantly lower than tone IA scores of normal articulating audios (t-test, *t(28)* = - 12.18, *p* = 1.1×10-12) and within the range of chance level (**Fig. 4c**). Therefore, behaviorally PA5 was not able to produce intelligible tonal speech during the task.

We then compared the tone-decoding accuracy of our framework on both participant PA5 and other normal articulating participant, the intended speech was consistently decoded, the average accuracy was 93%±3% and 95%±2% (mean [±SE]) with language model, 58%±9% and 51%± 2% (mean [±SE]) with language model, no significant difference was observed (t-test, with language model: *t(18)* = 0.48, *p* = 6.4×10-1, without language model: *t(18)* = −0.79, *p* = 4.4×10-1).

Finally, tone decoding accuracy and the tone IA scores of each syllable in the 4 articulating participants were significantly correlated (Pearson’s correlation *r* = 0.61, *p* = 2.9×10–5), indicating our tone decoder learned from behaviorally-relevant tone-related neural features, rather than other co-variants (**Fig. 4d**).

### Universal decoding framework with predefined hyperparameters is applicable on participants with great variations

ANN-based BCI decoders often encounter model hyperparameters such as depth and width of the network, kernel size and strides, dropout rate, etc. Training these models usually require optimization of these hyperparameters for each individual subject. A key factor of the generalizability of the BCI decoder is how robust the model performance is regarding different sets of hyper-parameters across participants. To get an overall understanding of heterogeneity across participants, we computed their best hyperparameter combinations of speech detector, tone decoder and syllable decoder chosen by the optimization process. These hyperparameters include smoothing window (***S***), the probability threshold value (***P_t_***), the off-time threshold (***T_off_***) and on-time threshold (***T_on_***), and the error permissive rate (***EPR***) which are related to the thresholding utterance onsets; filter lengths (***FL_ini_*** for the first convolutional layer***, FL_conv_*** for the rest convolutional layers, and ***L_pool_*** for max-pooling kernel) and strides (***ST_conv_*** for convolutional layers and ***ST_pool_*** for max-pooling layers) which determined the temporal feature of neuro-decoders; as well as depths (***C_layer_*** for sequential convolutional blocks and ***R_layer_*** for stacked recurrent layers) and widths (***C_dim_*** for number of filters for convolutional process, ***R_dim_*** for number of dimensions in each RNN process, ***D*** for dropout value) which determined the overall model architecture.. Very few optimized hyperparameter remained the same across all participants, while most optimal hyperparameters varied across participants with variations of age, gender, speech behaviors and electrodes coverage **(Fig. 5a-e)**. To figure out the impact of such heterogeneity on decoding performance in our framework, we picked the medium value of each hyperparameter and constructed a fixed pre-defined hyperparameter set. We then applied a universal decoder (UNI) on all the participants using this fixed pre-defined hyperparameter set. We found this universal decoder performed similar to (pair-wise t-test,PA1: *t(9)* = −0.62, *p* = 5.5×10-1, PA2: *t(9)* = 0.05, *p* = 9.6×10-1, PA3: *t(9)* = −2.00, *p* = 7.6×10-2, PA4: *t(9)* = 0.23, *p* = 8.3×10-1, PA5: *t(9)* = −0.55, *p* = 5.9×10-1) individually-optimized frameworks with hyperparameters optimized on individual participants (**Fig. 5f**). Also, the decoding performance of universal framework across participants was stable (oneway ANOVA, *F(4, 45)* = 1.11, *p* = 3.6×10-1). The UNI decoder even outperformed four control models only contained RNN and CNN part of the framework (CNN and RNN) with optimized hyperparameters. Mean WERs of decoding sentences through UNI remained lowest in all participants (**Fig. 5g**). Though the advantage was not significant in each individual participant due to relatively small sample size, it was significant after combining data of all the participants (paired t-test, UNI with CNN, *t(98)* = −3.29, *p* = 1.8×10-3, UNI with RNN, *t(98)* = −4.97, *p* = 8.3×10-6).

**Fig 5.**
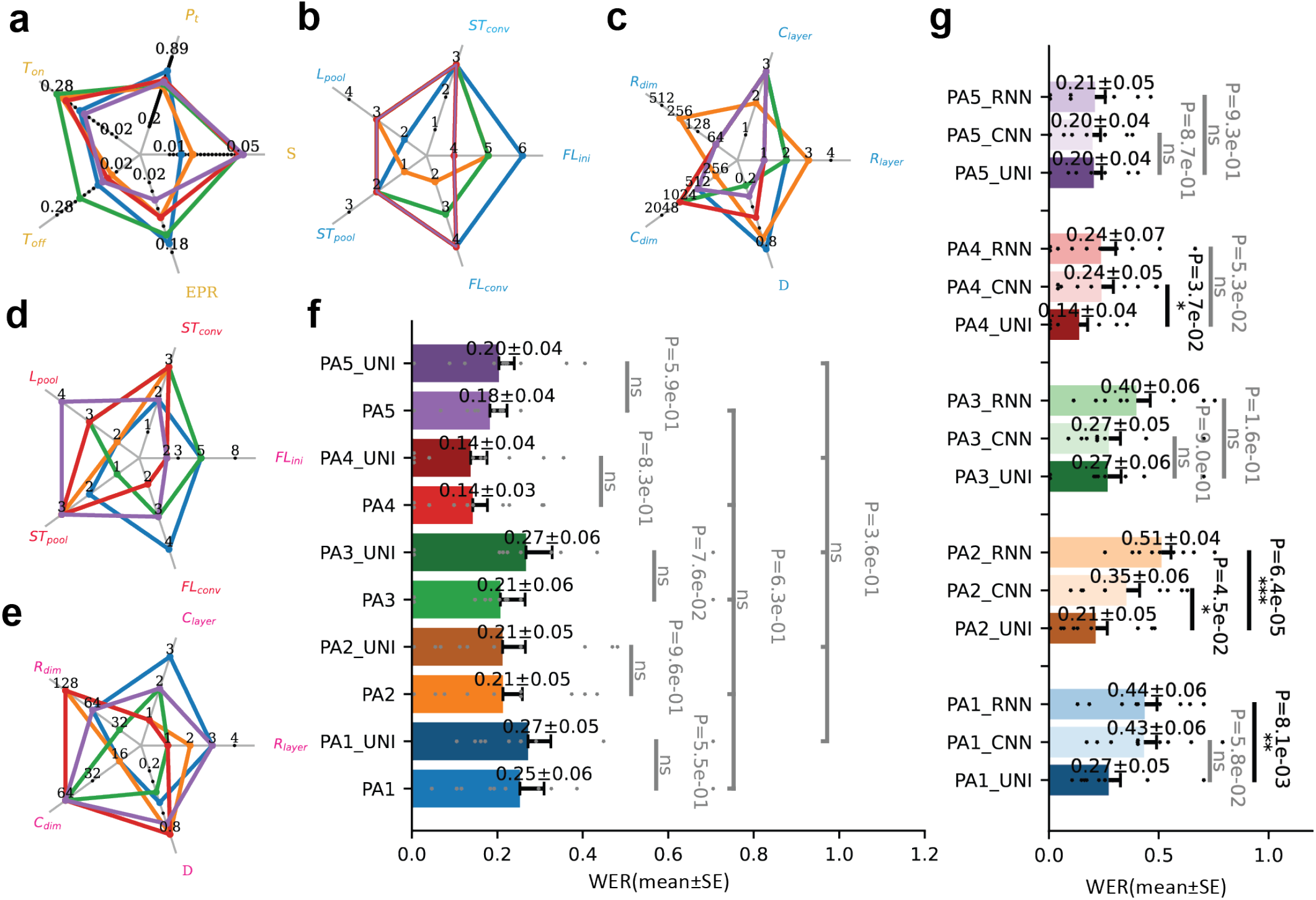
Evaluation of the neural decoder model hyperparameters across different participants. **a)** The optimal combinations of hyperparameters for speech detector in 5 different subjects (colors consistent in **f** and **g**): ***S*** represents the smoothing size, ***P_t_*** represents probability threshold, ***T_off_*** represents off-time threshold and ***T_on_*** for on-time threshold, ***EPR*** represents error permissive rate (detailed description of these hyperparameters see **Methods Section**. **b-c)** The optimal combinations of hyperparameters for the tone decoder: **b) *FL_ini_*** and ***FL_conv_*** represents the filter length of the initial convolutional layer and all following convolutional layers, ***ST_conv_*** represents for stride of all convolutional layers. ***L_pool_*** and ***ST_pool_*** represent for pooling length and stride of all pooling layers; **c) *R_layer_*** and ***R_dim_*** represent numbers of layers and hidden units of bidirectional Gated Recurrent Unit (GRU). ***C_layer_*** and ***C_dim_*** represent number of convolutional-pooling blocks and number of filters in each convolutional layer. ***D*** represents dropout value of each drop-out layers. **d-e)** The optimal combinations of hyperparameters for syllable decoder, similar to **b, c**. **f)** Comparison of decoding performances on each individual participant between individually-optimized decoders and the universal decoder with shared hyperparameters. In all the participants, universal decoder showed no significant difference from individually-optimized decoder (pair-wise t-test,PA1: *t(9)* = −0.62, *p* = 5.5×10-1, PA2: *t(9)* = 0.05, *p* = 9.6×10-1, PA3: *t(9)* = −2.00, *p* = 7.6×10-2, PA4: *t(9)* = 0.23, *p* = 8.3×10-1, PA5: *t(9)* = −0.55, *p* = 5.9×10-1). **g)** Comparison of decoding performances on each individual participant between universal frameworks (UNI) and customized control frameworks with hyperparameters optimized. In all the participants, UNI show lower WER than control models. In PA1, PA2 and PA4, these is significance difference (pair-wise t-test, PA1_UNI with PA1_RNN: *t(9)* = −3.38, *p* = 8.1×10-3, PA2_UNI with PA2_RNN: *t(9)* = −7.00, *p* = 6.4×10-5, PA2_UNI with PA2_CNN: *t(9)* = −2.33, *p* = 4.5×10-2, PA4_UNI with PA4_CNN: *t(9)* = −2.44, *p* = 3.7×10-2)between decoding performances of UNI and control models.

## Discussion

In this study, we present a brain-to-text framework capable of decoding natural tonal sentences from high-density ECoG recordings. We adopted a modular approach to delineate speech onset, base syllables, lexical tones, and leveraged contextual information through Bayesian likelihood and the Viterbi algorithm to enhance the decoding process. For natural speech, our proposed method achieved a tone decoding accuracy of 93%, similar to the behavioral performance of native speakers. The overall word error rate of decoded natural speech was as low as 14% in the best participant. Notably, we proposed a generalized spatiotemporal decoding framework for syllable and tone decoders. The robustness of our framework was evident across diverse participant profiles, including variations in gender, age, education, pronunciation behaviors, and electrode coverage, indicating that our model hyperparameters possess a high degree of generalizability. Significantly, the framework’s standardization of hyperparameters negates the need for extensive individual optimization, a step forward in practical application and scalability. Furthermore, our system adeptly managed the inherent language heterogeneity encountered in Mandarin, effectively handling the variances introduced by tone sandhi, regional dialects and individual speech patterns. Our decoder even showed robust performance in whispering conditions. These results underscore the potential for clinical applications in aiding patients with anarthria and broadening the communicative efficiency of BCIs.

This work extends the scope of ECoG-based speech brain-computer interface to natural speech of tonal languages. The past decade has witnessed significant breakthroughs in speech decoding and brain-computer interfaces using intracranial neural recordings. Previous works have also used ECoG to record local field potential directly from the sensorimotor cortex responsible for speech production. These works diverge in their decoding targets: some directly decode neural activity into speech sound or acoustics like spectrogram^2,26,27^; others map neural activity into discrete linguistic units such as words or phonemes^5,10,28^. Here we adopted a brain-to-text framework similar to the latter strategy where neural activity is first decoded into discrete syllables and lexical tones. Although directly decoding speech sound allows for continuous and infinite speech output, the quality of reconstructed speech is limited by the noisy neural signal. On the other hand, decoding into a finite set of syllables and tones extracts the invariant information from the noisy neural recordings. Furthermore, for cases like whispering or even completely covert speech, there is no ground truth of explicit speech output. As a result, brain-to-text may be feasible to such silent speech cases.

Our work underscores the importance of ventral sensory-motor cortex (vSMC) in speech production and decoding, particularly for tonal languages. Similar to our previous work^13^, we show that tonal speech production can be reliably decoded from neural activity in vSMC. Previous intracranial neurophysiology studies have investigated the spatiotemporal coding of the articulatory movements responsible of pitch control and phonetics^20–22^. A theoretical foundation of these studies is that there exist spatially distinct and distributed neural populations in the vSMC, representing different articulatory gestures corresponding to phoneme and pitch articulation. High-density ECoG recordings have proven to reliably cover the distributed network and dissociate these fine-grained neural coding^13,29^.

Our results demonstrate the excellent efficacy of our proposed light-weighted models in decoding articulatory movements, which is in line with a recent discovery that shallow feedforward networks achieve better performance on motor control than deeper and complex ones^30^. Articulatory movement is innervated via motor nerves which locates only one or two (considering inhibitory interneurons) synapses downstreaming the large pyramidal neurons (corticonuclear tract) originated from the motor cortex^31^. Based on intracranial neural recordings of higher signal-noise ratio (SNR), it is quite reasonable that light-weighted ANN is capable enough to replace the signal-processing function of such a few layers of synapses (including the lateral connections within the layers of cortex) ^32,33^. In our UNI frameworks, the numbers of trainable parameters in speech detectors, tone decoders and syllable decoders range from 2.3M to 6.7M. Such light-weighted frameworks not only showed less sensitivity to variations in hyperparameters, but also achieved better performance when trained on very limited amount of training data^34^. Such frameworks also reduce the responding-time and energy-consumption of computational infrastructures, which is a promising candidate for practical neuroprosthetic systems.

Our framework also provides insights into critical design considerations essential for speech BCI models. Previous studies of speech BCI typically relied on extensive hyperparameter optimization for individual participants who undergo chronic implantation^5,9^. Such decoding models may not be directly generalizable across different patients, resulting in repeated hyperparameters optimizing procedures for each individual. In contrast, our study reveals the feasibility of a universal, hyperparameter-optimization-free framework to five individuals, demonstrating its robustness across a spectrum of ages, genders, educational backgrounds, pronunciation habits, and variations in brain electrode coverage. Furthermore, as oppose to prior work where specific CNN and Recurrent Neural Network (RNN) models were designed for tone and syllable decoding respectively^13^, we proposed a unified CNN-RNN framework for both the tone and the syllable decoders in this study. Our discovery further validates that this combined CNN-RNN model achieved better decoding performance compared with baseline frameworks that employ only one of these network types. The benefits of our combined approach cannot be replicated through hyperparameter optimization alone. Our research highlights the potential for developing a broadly applicable, hyperparameter-independent framework for neural decoding. Although our universal framework has yielded stable performance across five distinct participants, future works remain to be done to consolidate its generalizability in patients with anarthria, across different Chinese dialects, and potentially other tonal languages.

This brain-to-text framework represents a pioneering effort in language BCI, designed to decode the full spectrum of Mandarin characters. Mandarin’s linguistic complexity is reflected in its use of over 6,000 commonly utilized characters, each a single syllable word. Yet, within this vast lexicon, there are only 416 unique segmental combinations of consonants and vowels. These unique combinations, combined with suprasegmental pitch features (4 different lexical tones), define 1664 unique tonal syllables^35^. Although there are some important phonetic features to the distinction standard 4 lexical tones such as turning point and ΔF0, amplitude, and speaker F0 range^36^. Other factors that determine the pitch contour in Chinese include the prosodic structure of the language, the interaction between syntax and phonology, and paralinguistic factors such as speech rate and tempo^18^. These factors can influence the pitch contour and contribute to the overall tonal patterns in Chinese, confusing native listeners listening to audio of each syllable clipped from natural sentences. Such bias obviously added challenges to speech BCI decoding tone exclusively from clips of neural signals while producing each syllable in natural sentences. The present decoding strategy inherently integrates the critical tonal features of Mandarin, effectively handling the variances introduced by tone sandhi, regional dialects and individual speech patterns, which are paramount for accurate speech communication, offering advantages over phoneme-level decoding. Besides, the current findings also highlight a pivotal insight: the decoding performance is not hindered by the length or complexity of the sentences, indicating the scalability of our framework for broader applications and more general settings in tonal language communication. Also, Mandarin characters are single syllable words usually consisted of only two or three phonemes, which is far less discriminative than most English. For example, "ji" versus "qi", or "zhi" versus "shi", as demonstrated in our study^2^. This structural intricacy of Mandarin, with its concise phonemic diversity, renders the phoneme-level decoding strategies used in recent speech BCI studies for non-tonal languages ineffective^7,8^. In response, our framework targets the decoding of Mandarin speech at the mono-syllable word level, aligning more closely with the language’s inherent structure and providing a practical blueprint for decoding its entirety—potentially expanding our current focus from the most frequently used 4×10 tonal syllables to all 4×416 tonal syllables.

In addition to the established use of high-density ECoG in speech decoding, various neural recording techniques offer distinct advantages in terms of coverage and temporal resolution. Examples include the Utah array^9,37^, stereoelectroencephalography (SEEG) ^38^, and neuropixels^39,40^. When choosing among these methods, a crucial consideration involves striking a balance between obtaining high-resolution neural signals, such as investigating fine-grained spiking properties of multiple single units or microcircuits underlying speech production within a limited area of the cortex^9,40^, and achieving broad coverage of cortical networks, such as collecting neural signals across the entire vSMC, which depicts a comprehensive view of neurodynamics of the functional regions^21,22,29^. Future works remain to be done to investigate the decoding capabilities of natural tonal languages using signals of varying coverage and resolution scales. This entails identifying the optimal trade-off point where high decoding performance aligns with decoding robustness across patients with high heterogeneity.

## Acknowledgements

Dr. Junfeng Lu is supported by STl 2030-Major Projects (2022ZD0212300) and the National Natural Science Foundation of China General Program (32371146). Dr. Yuanning Li is supported by the National Natural Science Foundation of China General Program (32371154) and Shanghai Pujiang Program (22PJ1410500). Dr. Jinsong Wu receives funding from Innovation Program of Shanghai Municipal Education Commission (2023ZKZD13) and National Social Science Fund of China Major Program (No. 22&ZD299).

## Competing interests

The authors report no competing interests.

## Data availability

Data relevant to this study are accessible from the authors under restricted access according to our clinical trial protocol, which enables us to share de-identified information with researchers from other institutions but prohibits us from making it publicly available. Access can be granted upon reasonable request. Any data provided must be kept confidential and cannot be shared with others unless approval is obtained. To protect the participants’ anonymity, any information that could identify him or her will not be part of the shared data. Source data and code to recreate the figures in the manuscript will be publicly released with code upon publication of the manuscript.

## Code availability

Code and source data to replicate the main findings of this study can be found on GitHub at https://github.com/yuanningli/tonal_BCI_decoding.

## Author contributions

J.L. and Y.Li. conceived and supervised the project. J.L., Y.Li., D.Z., Y.Liu., Z.Z., and S.L. designed the experiment. J.L, D.Z., Y.Q., and Z.Z. collected the data. D.Z., H.Z., X.H. finished the phonetic and phonological transcription. Y.Li., D.Z. designed the neural network. Y.Li., Z.W., D.Z., and W.L. designed the language model. D.Z. and Z.W. analyzed the data. J.L., Y.Li., L.C., K.X., and D.Z., interpreted the data. D.Z., Z.W., Y.Li., and J.L. wrote and revised the manuscript. All authors reviewed and approved the manuscript.

## Methods

### Participants

A total of five participants (a 41-year-old female, a 44-year-old female, a 54-year-old male, a 46-year-old male and a 30-year-old female) participated in this study. They were all patients with eloquent brain tumors who underwent awake surgery as part of normal clinical routine. Two 128 high-density electrode arrays were temporarily placed onto the lateral surface of the brain to collect the neural signals, and the participants were instructed to perform the speech tasks. All participants are native Mandarin speakers. An experienced neurosurgeon performed the grid placement, and the location of grid was determined based on the exposure and avoiding tumor.

The protocol was approved by the Huashan Hospital Institutional Review Board of Fudan University (HIRB, KY2019-538). All participants gave their written, informed consent prior to the surgery.

### Design of the sentence corpus

To have a representative set of phonological features in Mandarin and maximize the representativeness of our speech task, we selected the top 10 most frequently used open syllables with monophthong, which cover the pronunciation of nearly 25.9% of all Chinese characters^41^. Using these 10 syllables, we obtained 40 distinct Chinese characters with 4 lexical tones and constructed 29 Chinese words and phrases from these 40 characters. Finally, 10 sentences were constructed with these 29 phrases, which eventually consisted the sentence corpus used in our decoding task.

The 10 syllables chosen for this work is:

1. ’shi’, /ʂɚ/
2. ’de’, /tɤ/
3. ’ji’, /tɕi/
4. ’li’, /li/
5. ’bu’, /pu/
6. ’ge’, /kɤ/
7. ’qi’, /tɕʰi/
8. ’zhe’, /ʈʂɤ/
9. ’ta’, /tʰa/
10. ’zhi’ /ʈʂɚ/

The 40 Chinese characters used in this work and their corresponding tonal syllables (five-level tone marks):

1. 失: shi55
2. 十: shi35
3. 时: shi35
4. 实: shi35
5. 是: shi51
6. 的: de
7. 地: de
8. 得: de35
9. 机: ji55
10. 激: ji55
11. 极: ji35
12. 记: ji51
13. 计: ji51
14. 迹: ji51
15. 离: li35
16. 里: li214
17. 理: li214
18. 励: li51
19. 荔: li51
20. 不: bu35/bu51
21. 布: bu51
22. 哥: ge55
23. 搁: ge55
24. 个: ge51
25. 七: qi55
26. 奇: qi35
27. 其: qi35
28. 起: qi214
29. 契: qi51
30. 折: zhe35
31. 者: zhe214
32. 这: zhe51
33. 他: ta55
34. 塔: ta214
35. 织: zhi55
36. 枝: zhi55
37. 职: zhi35
38. 止: zhi214
39. 置: zhi51
40. 智: zhi51

*to simplify the decoding process, we use T1 (−) to denote 55, T2 (/) to denote 35, T3 (\/) to denote 214, and T4(\) to denote 51. The neutral tone syllables “的:de” and “地:de” were marked as T1 or T4 according to the actual pronunciation of each individual patient.

The 29-Chinese phrases used in this work (with translation in English):

1. 智者 sage
2. 记者 journalist
3. 荔枝 lichee
4. 七折起 thirty-precent off
5. 他 him/he
6. 他的 his
7. 他哥 his brother
8. 哥哥 elder brother
9. 不时地 often
10. 激励 encourage
11. 极其 extremely
12. 机智 smart
13. 织布机 loom
14. 这里的 here
15. 是 is
16. 的 DE (structure auxiliary)
17. 塔 pagoda
18. 不理 ignore
19. 是个 is a/an
20. 奇迹 miracle
21. 不计得失 do not care about gains and losses (Chinese Idiom)
22. 不得不 have to
23. 离职 resign
24. 搁置 shelve
25. 这个 this
26. 契机 opportunity
27. 其实 actually
28. 不止 more than
29. 七十 seventy

The 10 sentences used in this work:

1. 他哥不理他。

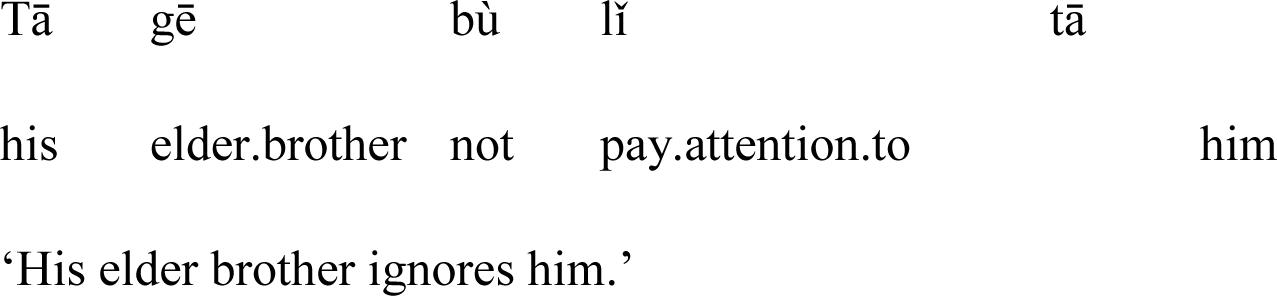
2. 他的荔枝七折起。

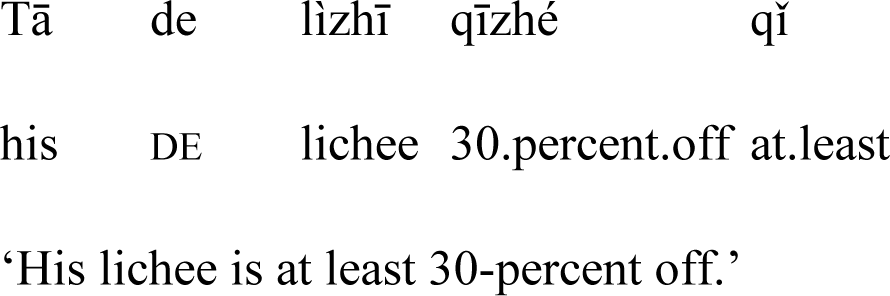
3. 记者不得不离职。

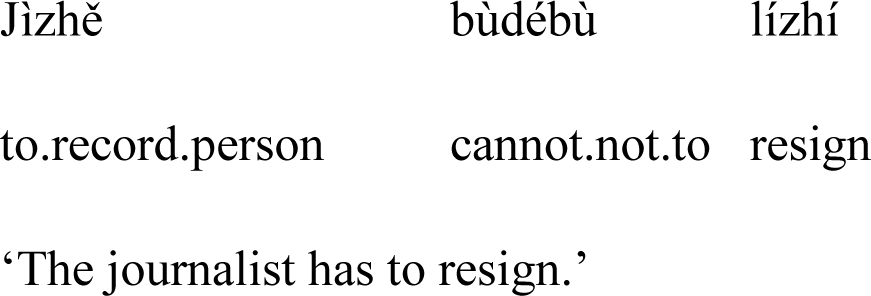
4. 他其实不止七十。

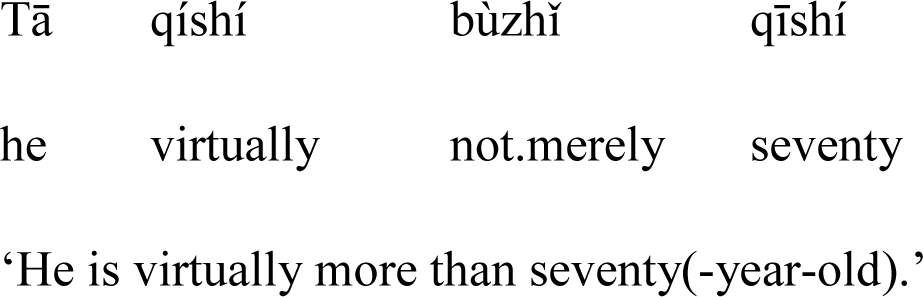
5. 哥哥不时地激励他。

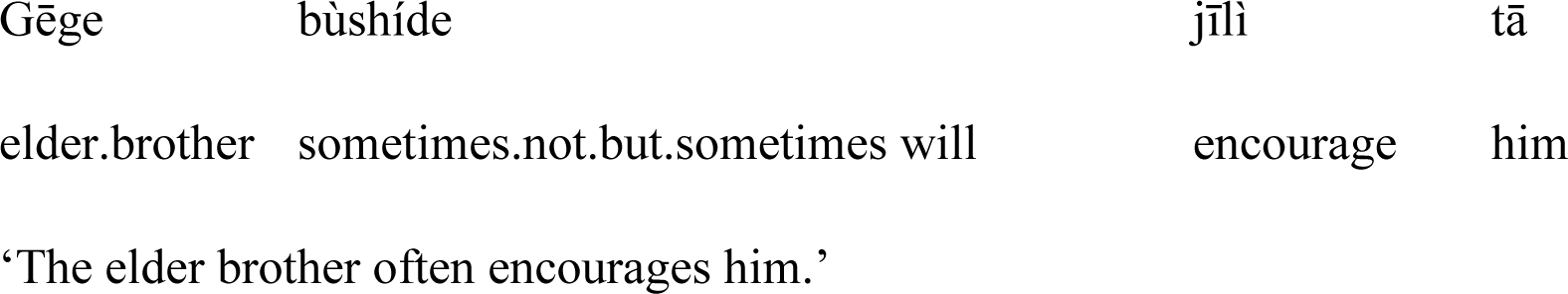
6. 他极其机智。

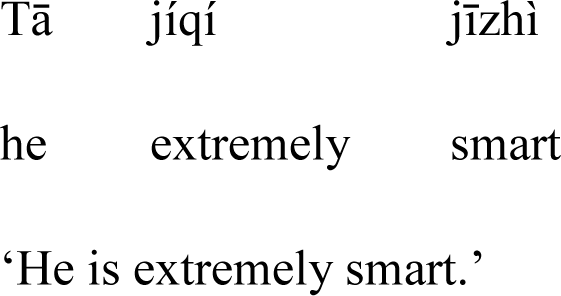
7. 织布机是这里的。

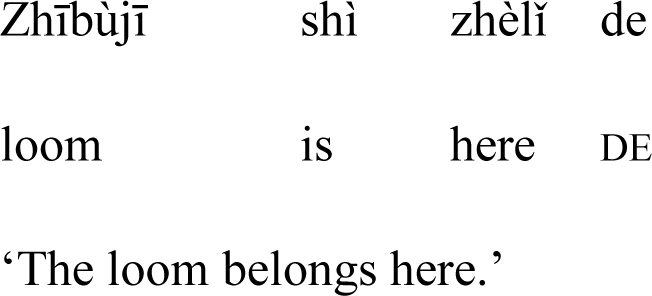
8. 这里的塔是个奇迹。

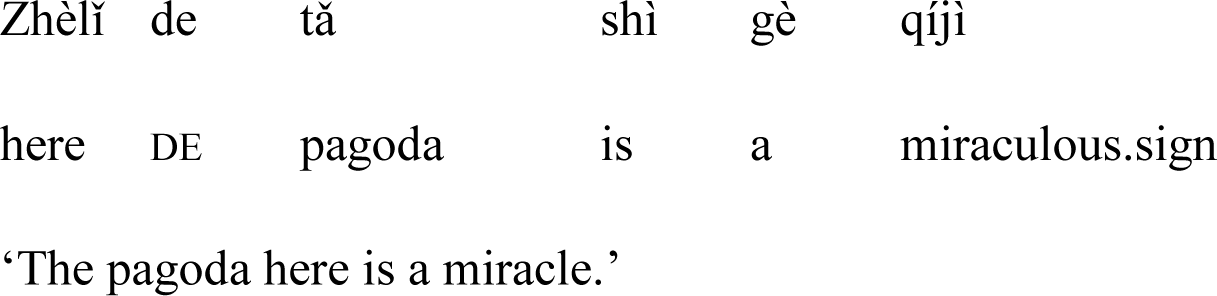
9. 智者不计得失。

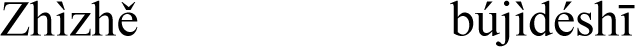
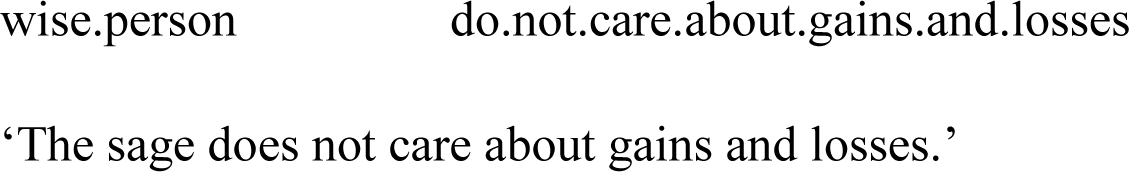
10. 搁置这个契机。

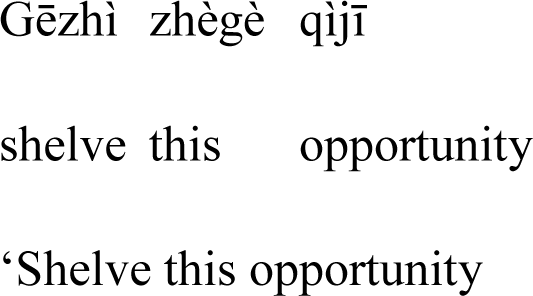

### Task design

Participants were guided by sequential visual cues to produce one of the 10 sentences consisted of 5-8 Chinese characters from a corpus of 10 base syllables and 40 Chinese characters with varied tones **(Fig. 1a)**. Within each trial, the participant was instructed to produce all 10 sentences once. These sentences were presented in a random order. Ideally, each participant produced 160 sentences (4 blocks × 4 trials/block × 10 sentences/trial), which yielded approximately 30 repetitions per tonal syllable. But not all participant completed the entire reading task (158-160 sentences were completed).

Each participant was guided by visual cues to perform block tasks. Each block started with a black cross at the center of white background on the screen, which lasted for 30 seconds. After that, the cross turned grey for 3 seconds, and one of the sentences in the sentence set was shown in the middle of the screen in grey text. With the individual Chinese characters turning black for 1.2 seconds in a sequential order from the beginning to the end of the sentence, the participant was instructed to pronounce the sentence in a relatively uniform speed following these go cues. The inter-sentence time interval and the inter-trial interval were both 3 seconds.

The first four participants (PA1-4) performed the speech task articulating, while the last participants (PA5) performed the speech task whispering. We synchronized audio recordings with ECoG recordings by utilizing a mounted microphone concurrently. We collected two types of blocks of the sentence task: first 3 optimization blocks (Trial 1-12, containing 118-120 sentences in total for each patient) and one evaluating block (Trial 13-16, containing 38-40 sentences for each patient).

### Data acquisition and signal processing

For each participant, two 128-channel electrode grids were placed by an experienced neurosurgeon. The anatomical positions to place the grids were chosen based on clinical exposure and avoidance of the tumor. During the tasks, electrocorticography and audio sound were simultaneously recorded using the Tucker-Davis Technologies ECoG system, at sampling rates of 3052 Hz and 24414 Hz, repectively. To exclude bad channels with artifacts or excessive noise, ECoG signals on each channel were visually and quantitatively inspected. High-gamma (70-150 Hz) frequency component was extracted via Hilbert transform after ECoG signals got down-sampled to 400 Hz^5,13^.

### Phonetic and phonological transcription

Transcriptions of the audio recordings, encompassing monosyllabic Chinese character, syllable, and tone labels, were manually annotated by a native speaker at the syllable level using Praat (Version 6.1.01, https://www.fon.hum.uva.nl/praat/) to ensure fidelity to the participants’ actual vocalizations ^5,13^. Unexpected voicing unrelated to language task (such as communication with clinicians) was excluded from samples used in training models.

### Computational modeling infrastructure

The training and testing of the decoding models were performed offline using cluster of multiple NVIDIA GPUs.

### Data splitting

We splitted the data for speech detector and the syllable/tone decoder model testing from electrode selection and hyperparameter searching. For responsive and discriminant electrode selection and hyperparameter optimization, only the optimization blocks were used, which contain Trial 1-12 with each trial consists of randomly arranged 10 sentences. The evaluation block contains remaining Trial 13-16.

During the optimization stage of the speech detector model, we used a six-fold nested cross validation (CV) and each fold consisted of 2 trials. At evaluation stage, we used a similar cross validation process, which leaves each trial in evaluation block for testing model, while the other 15 trials for training and validating. For speech detector models, we used 10% of non-testing data as validation set to perform early-stopping, while the left 90% for training, in each CV runs (**Fig. S1A&B**). For syllable/tone decoder model, we trained 5 (optimization stage) or 10 (evaluation stage) sub-models in each CV runs, with different potions of data performing early-stopping while left for training. The final model used for evaluation was an ensemble of 5 or 10 sub models. And the overall decoding performance was evaluated by averaging the test performance in all the CV runs (**Fig. S1C&D**).

### Speech-responsive electrodes

Speech-responsive electrodes were identified using two-sample t-test. Specifically, each time point in the [-400 ms, 800 ms] time window relative to the consonant onsets in each word was tested against the [−1800ms, −400ms] baseline time window before the onset of each sentence. If the results were significant (P < 0.01, Bonferroni corrected for the total number of electrodes and all times points) in 40 consecutive time points (100 ms), the electrode would eventually be marked as speech-responsive.

#### Tone discriminant electrodes

To pinpoint electrodes exhibiting discriminative characteristics among lexical tones, we aligned the high-gamma responses with the onsets of individual syllables again. Subsequently, we employed a one-way ANOVA to assess the potential differences in mean high-gamma responses across the four Mandarin tones. The time window for the average response spanned from −500 ms to 500 ms relative to the onset, encompassing a total of 400 time points. Significant time points were identified using a two-sided threshold of *P* < 0.05, with Bonferroni correction applied for both the total number of electrodes and time points^13^. To account for electrodes with multiple peaks of high-gamma activity discriminant for tone decoding, we set the following criterion: if there was a continuous 200ms time window in which more than half of the time points were significant, the electrode would be marked as tone-discriminant.

#### Syllable discriminant electrodes

Similar to tone discriminant electrodes, the syllable discriminant electrodes were defined using one-way ANOVA. We tested whether the mean high-gamma responses of the ten syllables were significantly different. The time window for the average response spanned from −500 ms to 500 ms relative to the onset, encompassing a total of 400 time points. The significant time points were determined via aforementioned criterion. Similarly, if there were a 200ms time window in which more than half of the time points were significant, the electrode would be marked as syllable-discriminant.

#### Cortical surface reconstruction and electrodes visualization

Electrodes on each individual brain were marked on preoperative T1 MRI in BrainLab® neuronavigation system, which were double-checked by a neurosurgeon via intra-operative photos^13^. The reconstruction of cerebral surface, anatomical labeling and plotting were performed via Freesurfer and customized python codes as previously reported^42^.

### Decoding framework

We built different neural network modules of speech detector, tone decoder and syllable decoder to decode speech onsets and offsets, syllable labels and tone labels from the neural activity, respectively. Inspired by previously published decoding models such as the EEGNet^43,44^, our decoders had separable CNN layers extracting within- and cross-electrodes spatiotemporal features, followed by GRU layers extracting sequential information. Finally, similar to previous brain-to-text works in non-tonal languages^5,9,10,37^, we also used a natural language model and a Viterbi decoder to combine the sequential outputs of the tone and syllable decoders and generate the entire sentence using maximum a posteriori probability **(Fig. 1b)**.

### Speech utterance detection model (speech detector)

The speech utterance detection model processed each time point of neural activity on speech responsive electrodes using the time window spanning from 0.25 second before the time point to 0.25 seconds after the timepoint. The speech/silent state of each time point was decoded based on the time-windowed neural activity.

We used the Torch 1.13.1+cu117 Python package to create and train the event classification model^45^. The event classification architecture was a CNN structure followed by a stack of three GRU layers with a latent dimension size of 256 and a dropout of 0.5 applied at each layer.

The initial CNN structure contains an 1D convolutional layer with a kernel size (***FL_ini_***) of 3 and a stride(***ST_conv_***) of 1 followed by a Leaky ReLU activation function to introduce non-linearity, and a max-pooling layer with kernel size (***L_pool_***) of 2 for temporal downsampling^46^. The initial processing, which involves convolution and pooling, enhances the model’s ability to efficiently extract local features from the input data. Subsequently, recurrent layers are incorporated to maintain an internal state over time and assimilate new individual time samples of input data, making them ideal for analyzing temporally dynamic processes. Additionally, the bidirectional nature of the Gated Recurrent Unit (GRU) enables the model to grasp both forward and backward temporal dependencies in the ECoG data, making it a fitting choice for event classification tasks requiring a comprehensive understanding of sequential context. Following the GRUs, we implemented a fully connected layer that projects the output of the last layer to probabilities associated with two target events: rest and speech. During training, the model is optimized to minimize weighted crossentrophy. The weighted crossentrophy loss was calculated using the ratio of the number of all the speech time points and the number of all the rest time points, to offset the bias caused by imbalanced samples. The batch size is 1024 and an Adam optimizer was used with a learning rate of 0.001. The training process stops after the validation loss no longer decreases for 10 epochs or after 50 epochs, ensuring that the model has sufficiently learned the underlying patterns in the data but has not yet overfitted. A schematic depiction of this architecture is given in **Fig. 2f**.

During testing, the neural network predicted probabilities for each speech event label (rest, speech) given the input neural data. To convert these predicted probabilities into the time stamps of speech onsets, thresholding of predicted speech probabilities is needed. A sliding window of smoothing size (***S***) was used to smooth the decoded probability timecourse. Next, a probability threshold (***P_t_***) was applied to smooth probabilities to binarized values (with a value of 1 for speech and 0 otherwise). We then sliced these binarized values by applying time thresholds for both a minimum continuous duration of silent period before the predicted onset, and a minimum duration of continuous speech period after the predicted onset (off-time threshold (***T_off_***) and on-time threshold (***T_on_***)). Detailed examples of thresholding process see **Fig. 3b**.

But the required continuous state is not strict, we set an error permissive rate (***EPR***) to allow a small portion of timepoints with incorrectly predicted speech event labels presented during both continuous silent period before the predicted onset as well as the speech period after the predicted onset. Besides, we will not predict two or more onsets within 0.5s, since normal speaking rate of our task do not allow such short interval and this update in algorithm speed up the iteration process.

In all, this process of obtaining onset events from the predicted probabilities was parameterized by five thresholding hyperparameters: the size of the smoothing window (***S***), the probability threshold value (***P_t_***), the off-time threshold (***T_off_***) and on-time threshold (***T_on_***), and the error permissive rate (***EPR***). Hyperparameter optimization was performed to eventually determine the exact values for these parameters for the evaluation stage.

### Tone decoder

For each speech utterance onset that was detected or manually aligned, tone decoder computed tone likelihood by processing the neural activity in tone-discriminant electrodes spanning from 0.2 second before to 0.6 seconds after the detected onset of speech.

The tone decoder consists of ten ensemble neural networks with the same hyper-parameters but different validation sets and training sets (see **Fig. S1D**). The input was high gamma ECoG data array shaped ***N* × *T***, ***N*** is the number of tone-discriminated electrodes, while ***T = 320,*** which is calculated by the time duration (0.8seconds) multiplies downsampled ECoG frequency (400 Hz). Each network among 10 networks consists of the following stages:

#### Stage 1

In the first stage, one-layer 2D convolution with the kernel size (***FL_ini_, N***) and a stride of (***ST_conv_*, 0**), followed by a batch normalization layer (momentum=0.1, affine=True, and eps=1e-5) applied to the output of the spatial convolution. After an Exponential Linear Unit (ELU) activation function. Max-pooling was then performed along the temporal axis with a kernel size of (***L_pool_,* 1**) and a stride of (***ST_pool_,* 1**). During training, we applied a dropout layer a dropout value of ***D*** in the end of each CNN block for regularization to prevent overfitting.

#### Stage 2

In the second stage, ***C_layer_*** convolutional pool blocks were applied after the aforementioned pooling layer in series, each consisting of a dropout layer, a convolutional layer with kernel size (***FL_conv_,* 1**) and a stride of (***ST_conv_,* 1**), batch normalization, ELU activation, and max-pooling identical with aforementioned ones. These blocks further processed and extracted features from the data. During the first two stages, the filter number was set equal to ***C_dim_***.

#### Stage 3

In the last stage, the output from the last convolutional pool block was fed into a stack of ***R_layer_*** bidirectional Gated Recurrent Unit (GRU) layers. Each GRU layer has ***R_dim_*** hidden units. The final hidden state of the last GRU layer was passed through a fully connected (dense) layer which project to 10 output units, representing the number of syllable classes for classification. **(Fig. 2g)**.

The model was optimized to minimize crossentrophy loss using a batch size of 8 and an Adam optimizer with a learning rate of 0.0005. The training process stoped after the validation loss no longer decreases for 50 epochs. The weighted crossentrophy loss was calculated in the similar same way as speech detector to offset the bias caused by imbalanced samples.

### Syllable decoder

For each speech onset that was detected, the syllable decoder computed syllable likelihood by processing the neural activity in syllable-discriminant electrodes spanning from 0.4 second before to 0.8 seconds after the detected onset of speech utterance. The structure and training process of syllable decoder was identical with the aforementional tone decoder, the only difference between them was the crossentrophy loss was not weighted, since the syllable is almost balanced-distributed in the task.

### Hyperparameter optimization and the universal framework

The speech detector, the tone decoder and the syllable decoder include a total of 25 hyperparameters. To find the optimal combination of these hyperparameters, we used the *hyperopt* Python package, which employs probabilistic sampling of hyperparameter combinations during optimization^5^. Across our experimentation, we utilized three distinct types of hyperparameter optimization procedures to fine-tune a total of 25 hyperparameters. (see **Fig. 5** for undefined hyperparameters, their searching ranges and their optimal values in each patient). During the optimization process, all the neural networks were tested through the six-fold (2 trials per fold) cross-validation.

#### Speech detection optimization

We used this procedure to optimize the size of the smoothing window (***S***), the probability threshold value (***P_t_***), the off-time threshold (***T_off_***) and on-time threshold (***T_on_***), and the error permissive rate (***EPR***). These hyperparameters were not related to the training of the ANN models. The predicted time points of speech onsets were derived from the available speech probabilities through the current combination of thresholding hyperparameters during each iteration of the optimization procedure. Subsequently, speech detection accuracy (***Acc***) was calculated to quantify the performance of the current combination of hyperparameter values. For timepoint of real onsets, we use 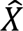 to represent the index of time points within 0.25s range of real onsets 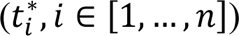. For time point of predicted onsets, we use 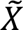 to represent the index of time points within 0.25s range of real onsets. Then, we calculated the ***Acc*** of 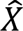 and 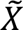. The formal expression of this objective function was illustrated through following equations:

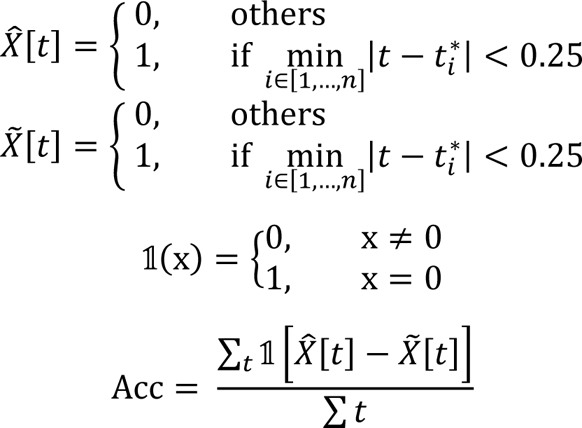

***Acc*** was calculated to qualify the performance of the speech detector with each hyperparameter value combination. During the optimization process of one model, 500 hyperparameter value combinations were evaluated.

Once the most optimized combination of hyperparameters was defined, we calculated the predicted onset time list for all the 12 trials via six-fold prediction, we compared them with the ground truth of onset of the 12 trials. If one onset time point in predicted list falls within 0.25s range of real onsets, the labels (including tone, syllable and Chinese characteristic) belongs to the manually aligned onset will be given to the predicted time point, which participated in the following hyperparameter optimization. For those time points cannot match with any real onset time points, no labels will be given and this part of the data will not participate in the following hyperparameter optimization. The similar match process will be performed in the evaluating stage, calculating confusion matrix for syllable, tone and word on the bases of onset detection. Because some real onset time points do not match with any detected onset time point, the number of syllables and tones that actually used in calculating confusion matrices of decoding performance after automated onset detection is less than the number of onsets detected.

#### Tone and Syllable decoder optimization

We used this procedure to optimize ten hyperparameters for both tone decoder and syllable decoder. These hyperparameters can be divided into two major groups according to their properties.

The first group includes initial convolution filter length (***FL_ini_***), stride of convolution (***ST_conv_***), filter length of following convolutional blocks (***FL_conv_***), max-pooling kernel length (***L_pool_***), max-pooling stride (***ST_pool_***), which are five hyperparameters related to the time dimension.

The second group includes the number of sequential convolutional blocks (***C_layer_***), number of layers of RNN (***R_layer_***), number of filters in each convolutional process (***C_dim_***), number of dimensions in each RNN process (***R_dim_***), and dropout value (***D***), which are five hyperparameters related to model architectures and sizes.

We performed model testing on manually-aligned-onset dataset and the dataset generated from predicted onsets by the previously optimized speech detector. And the final loss is the average value of loss on both datasets, in order to improve the model’s robustness when countering automatedly aligned neural signals in our final evaluation. Each optimization evaluated 500 different combinations of hyperparameter values. We used the same cross-entrophy loss functions to calculate the loss during the iteration.

#### The universal framework

The universal framework shares a predefined set of hyperparameters when testing on all 5 participants. We used the median value of each optimized hyperparameters among 5 participants as the predefined value of hyperparameters.

### Conversion from tonal syllables to Chinese words

Many Chinese characters are homophones that share the same pronunciations (including syllable and tone), which prevented us from directly generating Chinese sentences using decoded tonal syllables. To eventually decoding sentences consists of actual Chinese characters rather than sequences of tonal syllables, we designed a natural language model that computed the next-character transition probabilities given the previous Chinese character in a sequence. We first divided the sentences into the aforementioned 29 words and phrases consisting of 1 to 4 Chinese characters. We trained this model on a collection of sentences from the CCL corpus^47^ that included transferring pairs (transfers) between those words from the 29-word set. After that, we used a Viterbi decoder to determine the most likely sequence of words given the predicted tonal syllable probabilities from the tone decoder and syllable decoder, as well as the word-sequence probabilities from the natural language model^8^. With the incorporation of the language model, the Viterbi decoder was capable of projecting sets of probability (likelihoods) of tonal syllable to exact Chinese characters, which eventually decoded sentences from neural activity.

### Language model

#### Collection of the corpus

We used CCL corpus from Peking University^47^ to distill training dataset for our domain-specific language model. We first measure the number of transfers between each two phrases (the counts of the previous phrase transferred to the first word of the next phrase in the whole CCL corpus).

Due to the inequality of Chinese language, we set a cut-off of 512 counts, and then performed a ninth root for normalization. Since not all the transfers in our task appeared in CCL corpus, we also incorporate the transfer probability of our own task corpus into our language model.

#### Model fitting

We extracted all n-grams with n ∈ {1, 2, 3, 4} from each phrase in our task corpus. An n-gram refers a word lengthen n Chinese characters. For example, the n-grams (represented as tuples) extracted from the longest phrases “不计得失” in this approach would be:

1. (不)
2. (不计)
3. (不计得)
4. (不计得失)

Then we expanded our transfer probability matrix into n-grams. We added the number of transferring from *q*_*i*−1_ (such as (不计)) to *q_i_* (such as (不计得)). Except for full phrases, n-grams as partial components of phrases can only transfer from *q*_*i*−1_ to *q_i_*. The calculation of number of each transfering from a full phrase to the beginning of next phrases is aformentioned.

#### Hidden Markov Model (HMM) and Viterbi decoding

In this study, we employed a Hidden Markov Model (HMM) for neural activity and speech modeling similar to that used in previous research^5^. This model interprets neural activity within each time window at index *i* as observed states *y*_*i*_ and treats the word spoken during this period *w*_*i*_, along with its context ***c***_*i*_, as hidden states *q*_*i*_. The model assumed first-order Markov property and the current hidden state was fully characterized by *p*(*q*_*i*_|*q*_*i*−1_, *y*_*i*_). The transition probabilities, *p*(*q*_*i*_|*q*_*i*−1_), which represents the probability of transitioning from the n-gram at index *i* − 1 to the n-gram at index *i*, can be calculated by word counting from the corpus.

Viterbi decoder^5,48–50^ was used to identify the most probable sequence of hidden states from the observed neural activities. The algorithm used dynamic programming and considered the probabilities of transitioning between hidden states and the likelihoods of observed states. In the process of identifying the optimal hidden-state sequence, the algorithm calculated the probabilities of different potential sequences of hidden states. Each potential sequence, or Viterbi path, was defined by a unique sequence of hidden states (specific word sequence) and its corresponding probability based on the neural signals observed.

### Evaluation

#### Evaluation of independent decoding performances via Receiver Operating Characteristic (ROC) and Confusion Matrices

To evaluate the performance of our speech detector, we computed the Area Under the Curve (AUC) of the Receiver Operating Characteristic (ROC) curve for each participant. To evaluate the tone decoder and syllable decoder, we computed the classification accuracy and plotted the confusion matrix, based on both the predicted and the manually aligned speech onsets. We also calculated the classification accuracy and plotted confusion matrices of tonal-syllables and Chinese characters, on both the predicted and the manually aligned speech onsets, in order to further assess the capacity of the entire decoding system.

#### Evaluation of overall decoding performance via Word Error Rates (WERs)

To evaluate the overall performance of our neural-to-text decoder, we analyzed the decoded sentences using Word Error Rates (WERs) between the target and decoded sentences for each sentence. WER is a widely-used metric for evaluating predicted word sequences^5,51,52^. WERs of each decoded sentence in the evaluating blocks were calculated.

#### Comparison with published tone decoder

To further validate the complexity of tone of sentences compared with tone of single syllable, we applied our previously published monosyllabic tone decoder on tone decoding in sentence task. It is a sequential CNN-LSTM structure to generate the syllable label based on manually aligned neural activity when participants produce eight syllables "ma (tone1), ma (tone2), ma (tone3), ma (tone4), mi (tone1), mi (tone2), mi (tone3), mi (tone4)" ^13^. **(Fig. S2)**

#### Tone intelligibility assessment (IA)

Crowdsourcing-based listening tests are conventional for evaluating the quality of outputs in natural language processing^13^. To further validate the variance of lexical tones in natural sentences compared with the canonical tones of single syllables, we extracted the audio clips of each syllable utterance in the evaluation blocks of each participant, using the same timespan as the neural decoder. We shuffled the sequence clips by syllables to avoid any possible of leakage of semantic information. Evaluators were instructed to listen to the audio clip and then choose from the options (four lexical tones) which tone they had just heard^13^. For sentence tone IA, we extracted the audio clip of each sentence each patient pronounced in evaluation blocks. The evaluators listen to the audio of the entire sentence and then choose from the options for each syllable in the sentence. 20 evaluators (native Mandarin speakers) were instructed to listen to the isolated audio clips and to decide which tone they had just heard. For sentence tone IA, we extracted the audio clip of each sentence each patient pronounced in evaluation blocks. The evaluators were instructed to listen to the audio of the entire and then choose from the options which tone they had just heard for each syllable in the sentence. We recruited 20 native Mandarin speakers from the Medical School of Fudan University, which do not know the purpose of this test before the evaluation. The tone IA score was defined as the average accuracy of evaluators’ judgement.

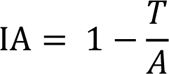

Where T is the number of error tone choices; A is the number of total tests^13^.

#### Comparison with baseline models

We evaluated the performance of our models and compared it against several baselines. The baseline models include (1) CNN models, which do not contain RNN structure, all the other parts remain identical with original optimized models. (2) RNN models, which do not contain CNN structure, all the other parts remain identical with original optimized models (**Fig. S3**). WERs of decoded sentences were calculated to evaluate the performance of these baselines.

#### Electrode contributions (saliences)

To quantify the contribution of each ECoG electrode to the brain-to-text decoding, we performed the electrode salience analysis^5,53^. In particular, we backpropagated the final loss function of each ANN module to each electrode in the input layer by computing the gradient. The magnitude of this gradient would quantify the amount of influence that a unit perturbation in each electrode would have on the final output of the neural network. We computed the Euclidean norm across time and across evaluation blocks of the resulting gradient values for each electrode. Finally, we normalized each set of electrode saliences by calculating their root mean square values **(Fig. S4)**.

## Supplementary Information

**Fig. S1.**
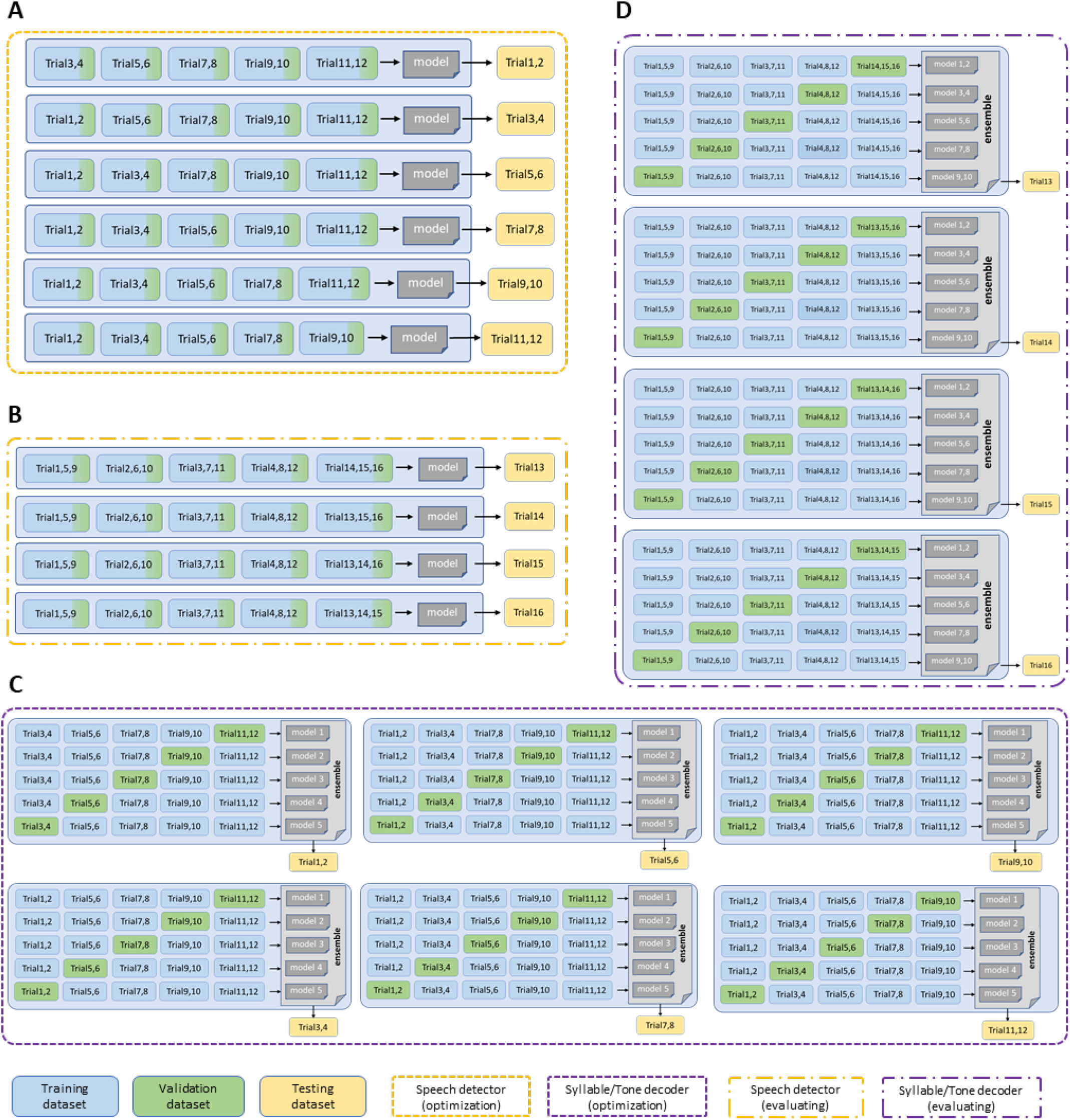
Schematic depiction of data organization. A. During the optimization stage of the speech detector model, we used a six-fold nested cross validation and each fold consisted of 2 trials, while 10% of data in each trial of the training set were separated as validation set in order to performing early-stopping. B. During the evaluation stage of the speech detector model, we used a similar cross validation. Still, 10% of data in each training trial were separated as validation set in order to performing early-stopping. C. During the optimization stage of the syllable/tone decoder model, we used a six-fold nested cross validation and each fold consisted of 2 trials. In each fold, we trained 5 sub-models with different potions of data performing early-stopping while left for training. The training/validation/test split was 4:1:1. D. During the evaluation stage of the syllable/tone decoder model, we used a similar cross validation process. In each fold, we trained 10 sub-models with different potions of data performing early-stopping while left for training. The training/validation/test split was 12:3:1.

**Fig. S2.**
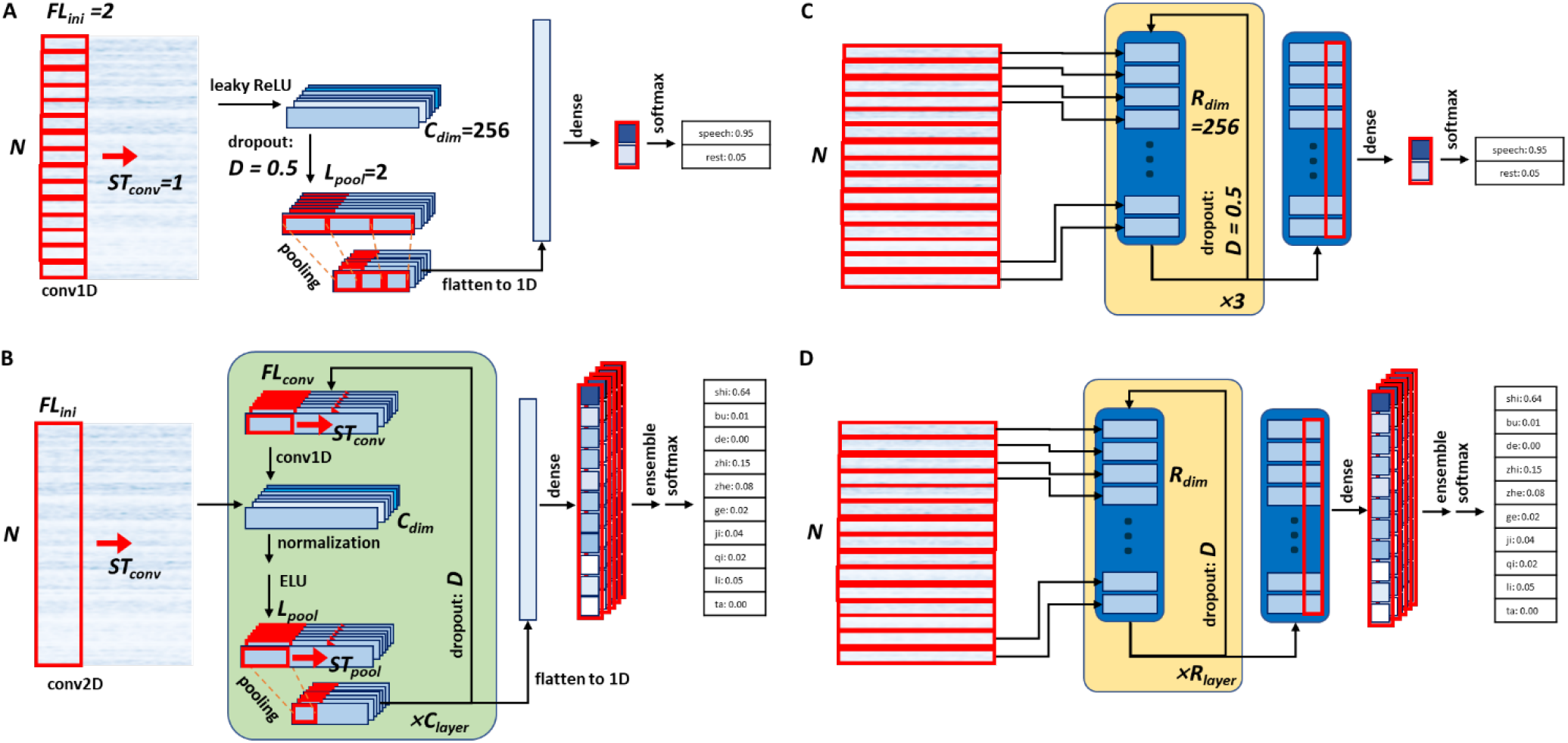
Schematic of control models for speech detector and tone/syllable decoder. A. CNN baseline for speech detector, which do not contain RNN structures, and all the other parts remain unchanged. B. CNN baseline for tone/syllable decoder, which do not contain RNN structures, and all the other parts, including optimized hyperparameters, remain unchanged. C. RNN baseline for speech detector, which do not contain CNN structures, and all the other parts remain unchanged. D. RNN baseline for tone/syllable decoder, which do not contain CNN structures, and all the other parts, including optimized hyperparameters, remain unchanged.

**Fig. S3.**
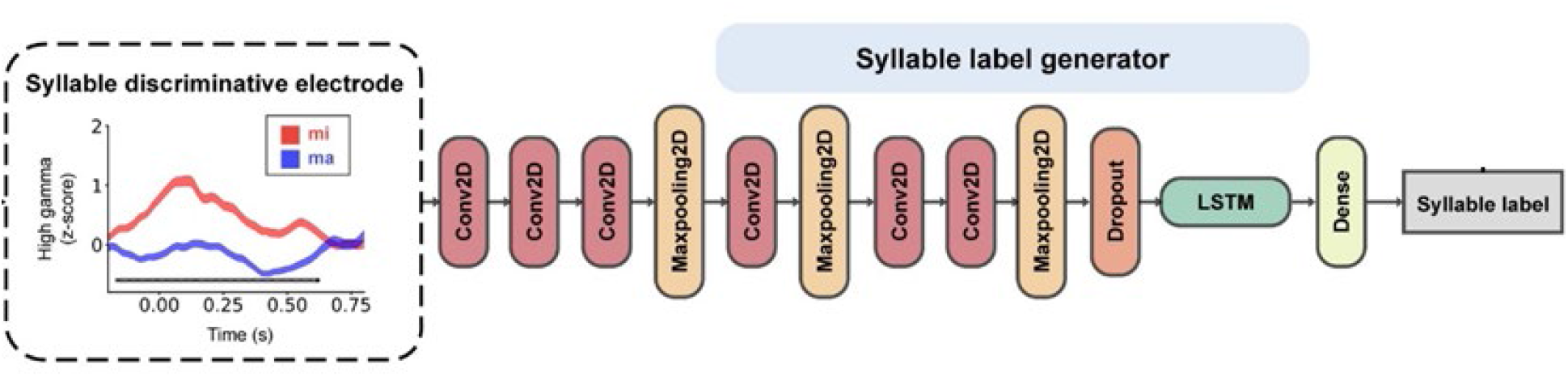
Schematic of our previously published tone decoder which successfully decoded tone from intracranial neural signals while participants articulate isolated monosyllables of four target tones. We applied our previously published monosyllabic tone decoder on tone decoding in sentence task. It is a sequential CNN-LSTM structure to generate the syllable label based on manually aligned neural activity when participants produce eight syllables "ma (tone1), ma (tone2), ma (tone3), ma (tone4), mi (tone1), mi (tone2), mi (tone3), mi (tone4)"^13^.

**Fig. S4.**
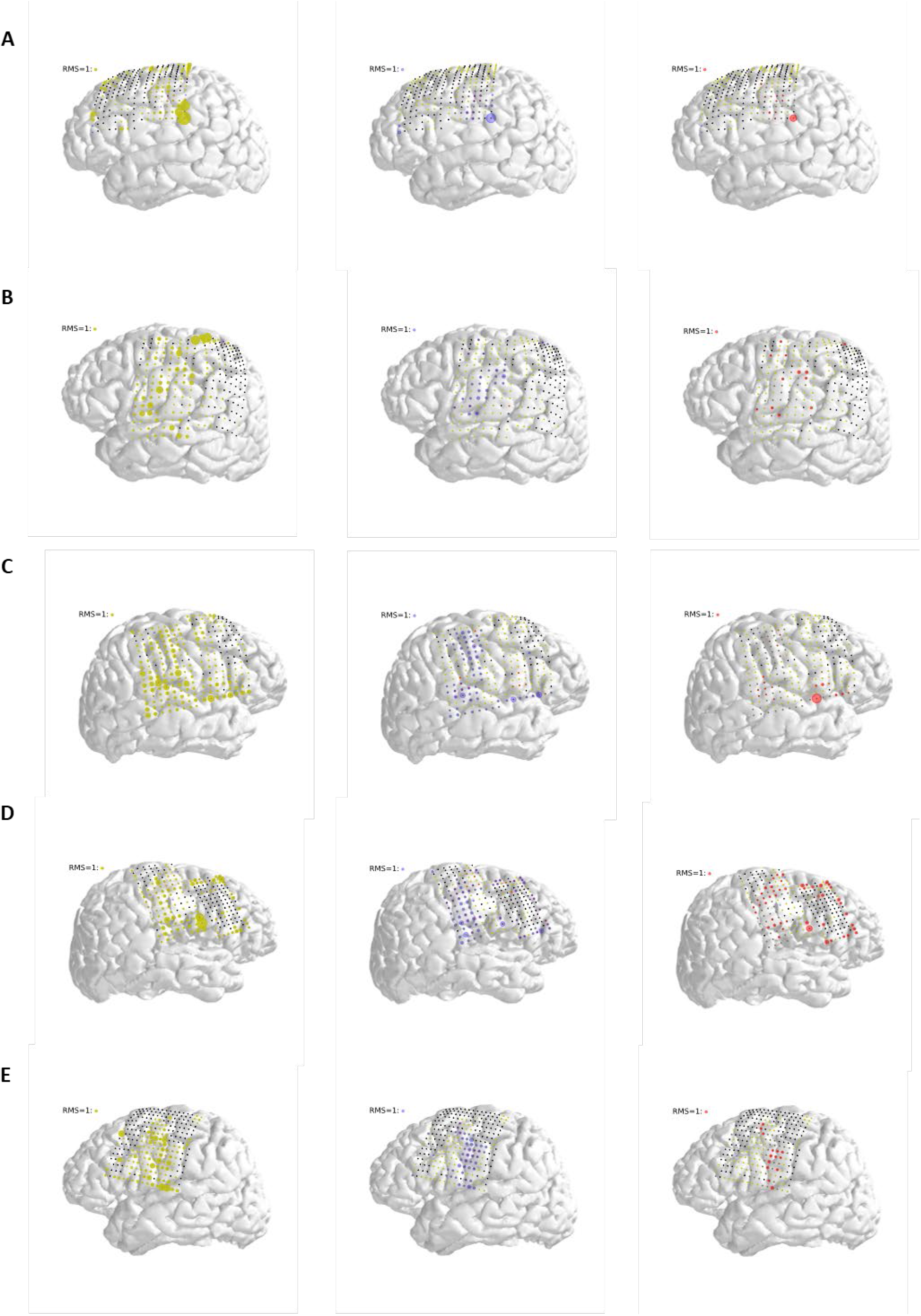
Participants’ brains reconstruction overlaid with the locations of the placed electrodes and their relative contribution of each selected electrodes in decoding onset/syllable and tone. A-E shown PA1, PA2, PA3, PA4 and PA5. Left part shown speech responsive electrodes in yellow shadows, middle shown syllable discriminative electrodes in blue shadows, while right shown tone discriminative electrodes in red shadows. Radius of shadows around electrodes shown contribution normalized root mean square (RMS), with a scale shown at the upper left of each subplot. The colors of the electrodes themselves represent whether they are speech responsive or syllable/tone discriminative, which is the same as **Fig. 2a-e**.

## Notes

### Competing Interest Statement

The authors have declared no competing interest.

### Summary of Updates

Methods section updated: descriptions of the speech detector, the language model, the Viterbi decoder, and the evaluation of electrode saliency updated. Reference updated.

